# Oxygen-responsive p53 tetramer-octamer switch controls cell fate

**DOI:** 10.1101/841668

**Authors:** Shashank Taxak, Uttam Pati

**Affiliations:** Transcriptional & Human Biology Laboratory, School of Biotechnology, Jawaharlal Nehru University, New Delhi-110067, India

**Author notes:** To whom correspondence should be addressed: **Uttam Pati:** Transcription & Human Biology Laboratory, School of Biotechnology, Jawaharlal Nehru University, New Delhi-110067, India.

**Keywords:** Oxygen sensor, Metastable, Switch, Chaperone, Prion

## Abstract

Solid tumors require an efficient decision-making mechanism to progress through a gradient of hypoxia. Here, we show that an oxygen-sensory p53 tetramer-octamer switch makes cell decision for survival or death in variable hypoxia. Trapping homo-oligomers from biosynthesis cycle, we found a metastable p53 tetramer in cells. Under the operation of switch, tetramer segregates the p53 character of a tumor suppressor and promoter. The p53 switch generates a pattern of its *on*-*off* state in time that is specific to the strength of hypoxia. A bidirectional tetramer-octamer conversion in *on* state decides the restoration of basal state by forward and programs apoptosis upon the reverse shift via p53-MDM2 loop. However, reversible dimer-tetramer transitions in *off* state trigger chaperoning of HIF-1 complex by tetramer in forward and oncogenic gain-of-function by prion-like dimers in reverse direction. Temporal *on*-*off* patterns calibrate stabilized p53 pool by defining the abundance of dimer, tetramer and octamer that ultimately decides diverse cellular outcomes in hypoxia. Through multi-chromophore FRET, we further show that chaperoning of HIF-1 may modulate angiogenesis through a possible flip-flop of the p53T-HIF-1 complex upon DNA. Our results demonstrate how p53 can sense oxygen and act upon its homo-oligomerization states to control cell fate in hypoxic tumors.

## INTRODUCTION

The role of p53 as a decision-making transcription factor in activating genes as part of specific gene expression in DNA damage repair, cell cycle arrest, apoptosis and senescence, has led to propose multiple mechanisms (1). p53 is inactive with loss of its apoptotic potential in the hypoxic region of solid tumors (2) and it remains unknown whether p53 transforms this state in order to control tumorigenesis through a gradient of oxygen. Variable tissue oxygenation in solid tumor triggers p53 conformational transitions from wild-type (WT) to mutant-like (ML) states, or *vice versa*, in order to control cancer progression (3) and it further modulates post-translational modifications in myocardial infarction dictating survival or death of the cell (4). Besides, change in ROS levels differentially controls p53 post-translational modifications in controlling p53-mediated apoptosis and necrosis (5). However, the mechanism of p53’s varied dynamics under oxygen stress is unknown. The cellular events operate through chemical switchboards, gene-protein loops and protein circuits in response to external stimuli. Molecular switches such as bacterial iron-cluster protein that senses molecular oxygen, iron, nitric oxide in countering to changing circumstances (6), GTP-binding between the GDP-bound *off* and the GTP-bound *on* state (7,8) and ion-mediated conformational switches and switchable lipids reversibly shifting between two or more stable states in response to stimuli and in microenvironment have been reported (9,10). Protein Tox, a decisive molecular switch for exhaustion mode of immune cells, activates a genetic program that alters immune function (11). Molecular switching is also implied by environmental stimuli that regulate the quiescence of radial neural stem cells (12). It was shown earlier that in resting cells p53 dimer is the most abundant oligomeric species which neither promotes cell growth or cell death until a shifting occurs towards monomers or tetramers in determining cell fate decisions (13) suggesting a probable switch mode that operate via p53’s homo-oligomer system upon external stimuli. We hypothesized such oxygen-dependent sensory switch in p53 homo-oligomerization pattern that might transform p53 state and control cell’s survival or death in regard to tumorigenesis under variable hypoxia. 1.5% of the human genome is transcriptionally responsive to hypoxia (14) and 60% of solid tumors exhibit hypoxic as well as anoxic areas throughout the tumor mass although the extent of hypoxia is independent of tumor stage, size, histology or grade (15). Hence, it might be possible that such an oxygen-responsive switch would integrate p53 with Hypoxia-inducible factor (HIF) and other hypoxic pathways such as chromatin remodeling, translation regulation and microRNA induction, in contributing to co-ordinated cellular response via oxygen sensing (16).

We have established that oxygen regulates the p53 oligomerization pattern via a tetramer⇀octamer switch that is both flexible and reversible. Metastable p53 tetramer responds to oxygen in a subtle way in directing the cell to undergo apoptosis, angiogenesis or tumorigenesis. In response to fluctuating oxygen levels in the cell, tetramer⇀octamer conversion as an active molecular switch could well segregate p53 dual function of prion and chaperone either to drive the cell into survival or death for tumorigenesis or maintenance of homeostasis.

## RESULTS

### Oxygen-sensing metastable p53 operates through a tetramer-octamer (T⇀O) switch

In order to prove the oxygen-sensitive switch in p53 homo-oligomerization system, it was first necessary to analyze p53 homo-oligomers status in the basal state of the cells. p53 monomers (M) dominantly exist as dimers (D) which can further assemble into tetramer (T) (17). Two units of T may also associate as octamer (O) (18, 19). p53 can further exhibit higher-order (H.O.) aggregation in the cell (19). Hence, to analyze p53 homo-oligomerization status, it was necessary that each form is simultaneously analyzed in the cell pool. Methods like MALDI-TOF and DLS were not suited for such analysis in the cell. Moreover, FCS (13) or BiFC (20) methods could fail for the simultaneous analysis of these forms in the cell pool. Therefore, based on previous report (19), BN-PAGE based approach was selected for the segregation and analysis of different p53 assembly states (including O and H.O). Under normal growth conditions, p53 usually appears in transient oscillation cycles (21). These oscillations may consist of a mixture of multiple homo-oligomerization states (13). During accumulation and degradation phases of these cycles, it is possible that different homo-oligomers undergo interconversion for the purpose of p53 restoration. Moreover, continuous accumulation or degradation would affect p53 homo-oligomerization differently than a stabilized pool. A stabilized pool is the prerequisite for the p53 mediated effects upon external stimuli. We hypothesized that the oxygen-sensitive switch could be observed in homo-oligomers system by stabilizing p53 oscillation cycles in the basal state of the cells. To stabilize p53 pool in the basal state, we inhibited cellular synthesis and degradation by CHX (cycloheximide) and MG132 interventions. CHX inhibits total cellular protein synthesis and MG132 is a proteasome inhibitor for cellular protein degradation (Fig 1A). Simultaneous inhibition of both the processes is a “trap” design in a sense that it separates homo-oligomers in transient oscillation cycles from p53’s synthesis and degradation pathways (Fig 1B).

**Figure1.**
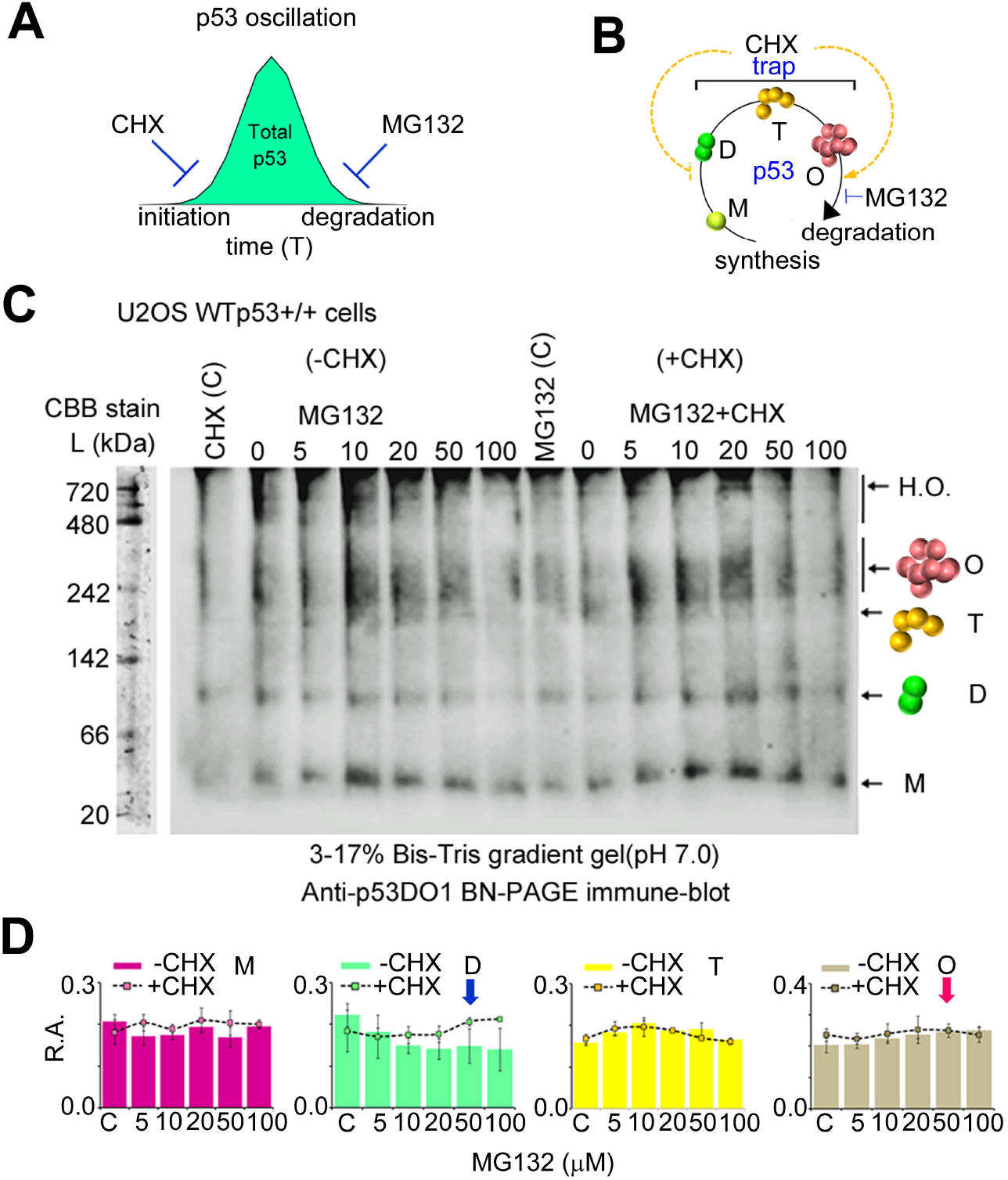
p53 tetramer exists as the metastable state in basal conditions. **(A)** Schematic representation of the homo-oligomerization trap generated by CHX (100μM) and MG132. **(B)** Spontaneous p53 oscillations captured by the trap in the basal state of cells. **(C)** Anti-p53 BN-PAGE immune blot shows p53 homo-oligomerization in basal state of U2OS cells by −CHX (only MG132 intervention) or +CHX (CHX+MG132 interventions) variants of the trap. 3-17% Bis-tris gradient gel (pH 7.0) shows p53 M, D, T, O and H.O. forms. O is observed as diffused smears. The immune density of O smear shows enhancement with an increase in MG132 dose (μM) in −CHX or +CHX variations. NativeMark protein standards were cut from the PVDF membrane after protein transfer and stained separately with coomassie brilliant blue G250 (CBB) dye. **(D)** R.A. calculation was performed by the densitometry of immune blots that identifies D↽T (blue arrow) and T⇀O (magenta arrow) conversion as an indicator of metastability of p53 T through −CHX and +CHX trap variants in the basal state of the cells. Immune blot shown in (C) is the best representation of the data in (D). Values and error bars in (D) represent mean and standard deviation from three independent replicates of the experiment respectively.

p53 resides in a stable steady state under normal growth conditions. Therefore, to obtain a switch-like behavior of homo-oligomers, a dose- rather than time-dependent intervention strategy was used. Under inhibited protein synthesis (+CHX variant of trap), a step-wise decrease in p53 degradation by increase in MG132 doses can detect fluctuation in M, D, T, O or H.O. level both due to association and dissociation. However, under continuous protein synthesis (−CHX variant), diminishing degradation can favor detection mostly due to association and hence, can be used as control to validate homo-oligomer fluctuations due to dissociation. Such a strategic use of cycloheximide (CHX) and MG132-based intervention in the basal state cells followed by p53 segregation in 3-17% Bis-tris gradient gel through anti-p53 BN-PAGE immune-blotting allowed detection of p53 M (53 kDa), D (106 kDa), T (212 kDa), O (424 kDa) or H.O. (>424 kDa) and their authentication based upon molecular weight (M.W.) by Native Mark protein standards (19) (Fig. 1C). The fluctuation in the homo-oligomers level due to association or dissociation in −CHX or +CHX trap variant was quantified through R.A., a unit normalized quantity called relative abundance. The quantity is independent of total p53 level (and hence, GAPDH normalization is optional) which justifies fluctuations due to association or dissociation of p53’s assembled products under the trap conditions. When compared in +CHX and −CHX variants of the trap, R.A. detected the conversion of T into D (D↽T shift) by dissociation & O (T⇀O shift) by association that satisfies metastability of T in spontaneous p53 cycles (Fig.1D, magenta and blue arrows). It should be noted that in a resting cell population, a few cells generate p53 oscillation cycles in an asynchronous manner which leads to capture of very low amount of its total pool and little fluctuations with faint density of immune bands by the trap (also resulting in comparatively low quality of immune blot images). In such a case, HRP chemiluminescence from immune blots was captured by extended exposure to camera. In this process, to check whether blot images remained unsaturated for p53 species, pixel-wise distribution of chemiluminescence intensity was used to calculate variance and mean for each immune band and variance v/s mean plot was fitted with straight line. A linear trend justified unsaturated immune bands, a prerequisite for the R.A. determination by densitometry measurements of BN-PAGE immune blots (Fig. S1A), and authenticated trends in R.A. by −CHX or +CHX based traps at low p53 level. The metastability of T suggests a switch-like behavior of p53 homo-oligomerization. Owing to such property, p53 may switch to D↽T or T⇀O states upon stress. Surprisingly, in a 3-17% gradient, O migrated as diffused smear in close proximity with T. To analyze whether such smeary pattern is the result of modification of O in the basal state of the cell, first, BN-PAGE gradient was changed to either 6-12% or 5-15% in order to allow sufficient resolution of T and O and second, an exogenous p53 system was utilized due to very low level of endogenous p53 in basal state. O in normoxia (21% O_2_ or basal state) was compared with hypoxia (1% O_2_) or re-oxygenation (48h) stress conditions. 6-12% gradient showed diffused O smears in normoxia that gradually diminished in hypoxia or re-oxygenation (Fig. S1B). This indicates that O can exist as an additional modified form in the basal state of the cell. The decrease in immune density of modified O upon p53 degradation at higher doses of cisplatin (22) (Fig. S1C) and enhancement at higher MG132 dose under +CHX or −CHX variant of the trap in endogenous systems (see Fig 1B) indicates that they are ubiquitinated species. Additionally, the lower R.A. in +CHX as compared to −CHX setup suggests H.O. species participation in p53 degradation (Fig. S1D). Hence observed H.O. state might correspond to polyubiquitinated O in BN-PAGE immune blot.

Next, in order to prove the operation of metastable T via D↽T or T⇀O shifts in time- and oxygen-dependent manner, a similar CHX based trap was utilized at different oxygen (O_2_) concentrations (5, 1 and 0.1%). In this case, MG132 was not required as low oxygen stimulus ultimately induces similar effect i.e. p53 stabilization by proteasomal inhibition (23) (Fig.2A) which may vary in the rate as per oxygen crisis (see Fig 3A). Application of MG132 would not provide solely an oxygen-dependent stabilization profile of p53 by allowing accumulation of its degradable fraction. As degradable fraction may follow D⇀T⇀O path, it would hinder the differentiation of D↽T and T⇀O shifts based on fluctuation in homo-oligomers R.A. at different oxygen conditions by BN-PAGE. As H.O. species were degradable fraction and not essential for our analysis, 3-17% was changed to 5-15% Bis-tris gradient gel in order to sufficiently resolve M, D, T and O (Fig.2B, C). Anti-p53 immune blotting did not show the diffused O under hypoxic conditions by CHX trap confirming that ubiquitinated or degradable fraction was completely eliminated from the pool and hence, homo-oligomers signals explicitly constituted dynamics of metastable T in oxygen- and time-dependent manner. 6h CHX treatment (T2) at any time in hypoxia (T1) was sufficient to completely eliminate the degradable p53 fraction of cell pool and as a result, durations T2>6h have no significant effect over net stabilized p53 level (compare +CHX signals at T1=18 and 24h or 42 and 48h for T2=6 and 24h in Fig. 3A or SDS PAGE immune blots in Fig.5A, B). Hence, a decreasing pattern of T2 for increasing T1 was designed to simultaneously process 0-72h T1 samples for the analysis by BN-PAGE. M appeared at insignificant level in the pool which could be the result of its rapid co-translational association into D (17) and hence, it was not considered for R.A. calculation. 5% O_2_, which lies in the range of normoxia in tissues (24), maintained a steady-state level of T via D⇌T⇌O during 0-72h interval (Fig. 2D, left panel, also see Fig 5B, upper panel). However, at hypoxic levels, D⇌T⇌O state is perturbed to either **D**⇌**T**↽O at 1% or D⇀**T**⇌**O** shifts at 0.1% O_2_, indicating direct oxygen sensing via metastable p53 T (Fig. 2D, magenta arrows in middle and right panel).

**Figure2.**
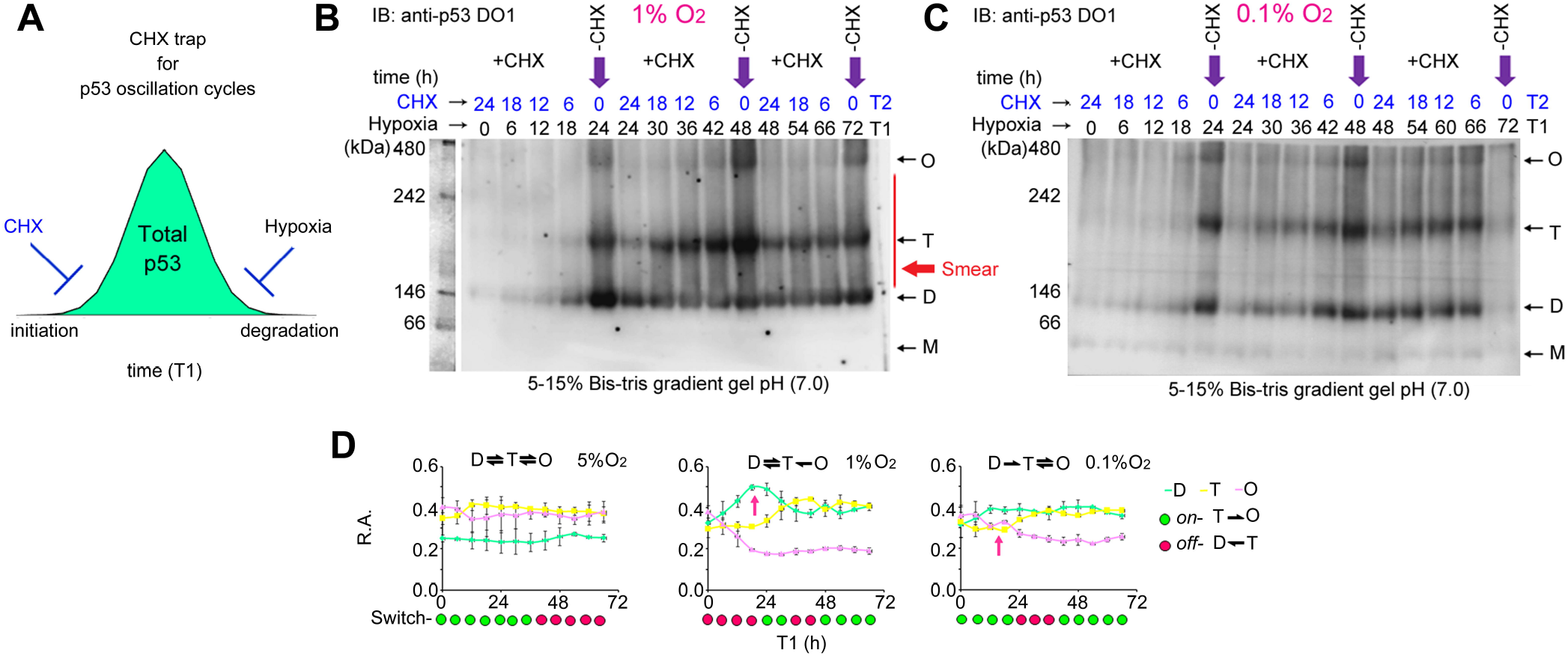
Metastable p53 T operates via an oxygen-sensitive T⇀O switch. **(A)** Schematic representation of the CHX trap in a hypoxia gradient. **(B, C)** To determine metastable p53 T dynamics in response to hypoxia, CHX trap design in (A) was used to capture p53 homo-oligomerization dynamics by anti-p53 BN-PAGE immune blotting at 1, 0.1 or 5% O_2_ (immune blot is shown in Fig. 5B). To sufficiently resolve each homo-oligomer (especially T and O) 5-15% Bis-tris gradient gel (pH 7.0) was utilized. T1 represents duration for which HCT116 p53+/+ cells were exposed to hypoxia before CHX treatment. Purple arrows indicate p53 pool segregated in its constituent homo-oligomers without CHX trap. T2 represents the duration of CHX for hypoxic cells. 24h>T2>6h was always maintained for p53T dynamics in 0-72h T1. A red arrow in (B) shows p53 aggregating smears. Native protein standards were run in the same gel and after transfer of samples on PVDF membrane; its lane was cut and stained separately with coomassie brilliant blue G250. Due to inclusion of protein standards in 15 well gel, 60^th^ h sample for 1% O_2_ was analyzed separately or from other replicates. SDS-PAGE based analysis of total p53 pool and GAPDH loading control of immune blots in (B, C) is shown in Fig 5A, B or Fig S3F. **(D)** R.A. measurements from (B, C) show oxygen-sensitive p53T via shifts in equilibrium state (5% O_2_). Green and magenta circles correspond to *on-off* pattern of p53 switch deciphered at 6h. The magenta arrow shows enhanced dimerization or octamerization via T during initial durations that initiates shifts at 1 and 0.1% O_2_ respectively. Values and error bars in correspond to mean and standard deviation from three independent replicates of the experiment respectively and are best represented by the immune blots in (B, C) or Fig. 5B.

**Figure3.**
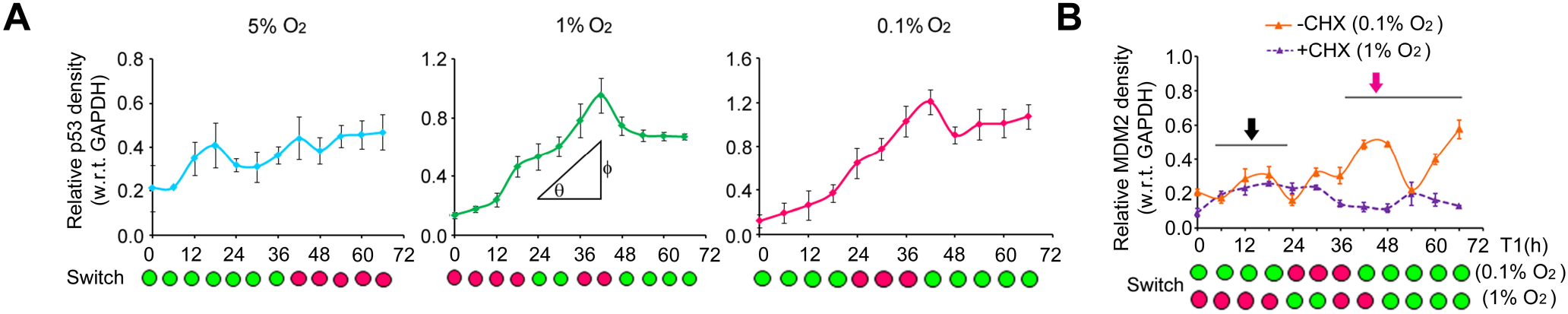
The function of T⇀O switch in *on* mode. **(A)** Rate of p53 accumulation by SDS-PAGE immune-blotting method using CHX-trap at 0.1, 1 and 5% O_2_. **(B)** MDM2 dynamics by SDS-PAGE immune blotting without or with CHX-trap at 0.1 and 1% O_2_ respectively. Data represent mean and error bars show s.d. from three independent replicates. Representative immune blots of plots in (A) or (B) are shown in Fig. 5A, B, Fig S3F or Fig S2A, B. Due to the inclusion of protein standards in 15 well gel, 60^th^ h sample for 1% O_2_ were analyzed separately or from other replicates.

D↽T step was observed exclusively in the presence of CHX under basal conditions, which complies with the fact that D⇌T inter-conversion depends upon post-translational modifications (14) independent of oxygen. On the contrary, T⇀O step showed variability in hypoxia gradient suggesting its oxygen-sensitive behavior. Hence, one could assume that T⇀O step might act as a switch in controlling the dynamics of metastable p53T in hypoxia. To demonstrate if metastable p53 operates through T⇀O switch, change in the rate of dissociation or association of T was compared at 5, 1 or 0.1% O_2_. BN-PAGE signals from the trap during 0-24, 24-48 and 48-72h intervals were first normalized with their −CHX variants at 24, 48 and 72h respectively (see Fig. 2B, C, purple arrows) and their log2(normalized value) or log_2_(fold change) were calculated and compared with the basal state level (broken line) at each interval (Fig. S1E). The measurements provided dissociation (D↽T) and association (T⇀O) rate of T relative to the basal state (0). From these measurements, we designated an enhanced rate of D accumulation via D↽T as *off* (magenta circles) and enhanced rate of O via T⇀O as *on* state (green circles) of the T⇀O switch. At 5% O2, O was found to be mostly eliminated from the pool. Hence, we designated *on* state during 0-36h interval based on minimal accumulation rate of D and T that would result from a continuous D⇀T⇀O→degradation step in the normoxia conditions to restore p53 pool. However, the rise in D and T levels with minimal O in the p53 pool can only result from *off* state of T⇀O in post 36 h interval. A closer examination revealed that *off* state of the switch directs D⇌T⇌O towards D⇌T↽O whereas *on* status allows D⇀T⇌O shift in time and oxygen-dependent manner thus indicating its function as an oxygen-sensory module for metastable p53 in hypoxia gradient. It is notable that the *on* status in late durations (48-66h) maintains a stable O level by balancing T↽O via T⇀O at 1% O_2_ in contrast to 0.1% O_2_ where T⇀O overcomes T↽O to enhance O level (Fig. S1F).

### An on mode tetramer ⇌ octamer state controls p53-MDM2 loop and apoptosis

It seems that the extent of oxygen crisis calibrates the duration of *on-* and *off* modes of T⇀O switch in terms of the proportion of D, T and O in the stabilized p53 pool. Besides, rate of stabilization (θ), as observed from anti-p53 SDS-PAGE immune blots, reveals that switch may control p53 degradation and accumulation by duration of its *on* and *off* modes at 5, 1 or 0.1% O_2_ respectively (Fig. 3A, SDS-PAGE immune blots are shown in Fig. 5A,B and Fig. S3F). However, the stabilized p53 may differ by its net amount (ϕ, y-axis) at variable oxygen due to the different proportion of its higher-order homo-oligomers in the total pool. O is most likely to get utilized for rapid degradation of p53 as it was found mostly in the ubiquitinated state under normal conditions. By contrast, D would be preferred for the enhanced accumulation as it possesses a higher stabilization tendency (13). **T**⇌**O** controls p53 octamerization by D⇀**T**⇌**O** shift and hence, can induce both p53 restoration and activation via p53-MDM2 loop in *on* mode. Anti-MDM2 SDS-PAGE immune blot showed the rate of MDM2 accumulation is enhanced in *on* when compared to *off* mode (Fig. 3B, Fig. S2A, B). However, abrupt fluctuations encountered in its rate might be the result of an additional p53-independent MDM2 post-transcriptional down-regulation effect in hypoxia (23). It should be noted that CHX-trap was essential at 1% whereas it was optional at 0.1% O_2_ in order to obtain actual MDM2 dynamics due to **D**⇌**T**↽O and D⇀**T**⇌**O** shifts during *off* and *on* modes of the switch (compare with Fig. 1D). Two modes maintain a basal MDM2 level during initial intervals (0-24h) (black arrow) in which a sufficient amount of T is still unavailable at both 0.1 and 1% O_2_ mostly via D⇀T⇀**O** and **D**↽T↽O respectively (compare with Fig S1F). This means that both the modes can achieve the required amount of T for MDM2 accumulation specifically via **T**↽O step in late intervals (42-66h, magenta arrow). Although **T**↽O seems functional for p53 activation by the assistance from D⇀**T** in *on*, it is deactivated by **D**↽T in *off* mode of switch, suggesting p53-MDM2 loop activation is associated exclusively with *on* mode D⇀**T**⇌**O** shift. Unlike 5%, 0.1% O_2_ shows a relatively higher stabilization of p53. **T**⇌**O** achieved at higher p53 level signifies an inefficient MDM2 activity that could be the result of uncoupling of p53 and MDM2 in an oxygen-dependent manner (25). In such circumstances, abrogated p53-MDM2 loop might result in terminal cell fate (26) in *on* mode via T⇌O. To check this, apoptosis was compared in HCT116 p53+/+ or HCT116 p53−/− cells by flow cytometry-based assays at 5, 1 or 0.1% O_2_. Exposed cells were trypsinized and cell suspensions were stained with membrane-impermeable YO-PRO1 and Propidium iodide (PI) fluorescence dyes. YO-PRO1 and PI differentially stain apoptotic and necrotic fractions based on the extent of their plasma membrane disintegration which causes variability in the internalization of these dyes by the cell. A two-dimensional plot of YO-PRO1 v/s PI fluorescence intensity identified apoptosis through high YO-PRO1 & low PI, necrosis by low YO-PRO1 & high PI and viability by low YO-PRO1& low PI fraction of cells. HCT116 p53+/+ cells showed approx. a two-fold increase in apoptosis at 0.1% (17%), almost similar to re-oxygenation (23%), as compared to 1% O_2_ (4%) confirming the role of *on* mode in apoptosis through **T**⇌**O**. By contrast a two-fold decrease in apoptotic fraction at 1% (4%) as compared to 5% O_2_ (8%) in HCT116 p53+/+ and the observed low apoptosis (3%) in HCT116 p53−/− at 0.1% O_2_ indicates the role of *off* mode in survival through **D**⇌**T** (Fig. S2C). Similar results were obtained in another variation of apoptosis assay using YO-PRO1 v/s FSC (forward scatter) analysis. FSC depicts the size of the cells which is usually reduced during apoptosis. In this case, a high or low YO-PRO1 cell fraction at lower FSC (than live controls) detects late (L.A.) or early apoptosis (E.A.) stages in hypoxia gradient (Fig. S2D). Gene-expression analysis by qPCR and micro-array at less than 0.1% O_2_ has shown activation of p53 downstream genes responsible for apoptosis (27) and microarray analysis of p53-dependent gene expression at 1% O_2_ has shown that the transcriptional response by p53 favors the cell cycle arrest over apoptosis (28). In such a case, our finding strongly links an on mode T⇌O shift with transcriptional activation of p53 for apoptosis.

### p53 tetramer physically interacts with HIF-1 complex

To check if T⇀O switch is responsible for linking to HIF-1 for efficient oxygen sensing, different HIF-1α complexes were analyzed in hypoxic HCT116 p53+/+ or HCT116 p53−/− cells by anti-HIF-1α BN-PAGE immune blotting (Fig. 4A). Phosphorylated and non-phosphorylated HIF-1α were identifiable at 120 kDa (29) by 3-15% Bist-tris gradient gel (purple arrows). Interestingly, a higher-order HIF-1α complex (242kDa<M.W.<480 kDa, blue arrow) sequestered HIF-1 (M.W.≈212 kDa, yellow arrow) in HCT116 p53+/+ cells. This provides a possibility of p53 and HIF-1 complex. Besides, higher-order aggregates of HIF-1α (M.W.>480 kDa) appeared specifically in HCT116 p53+/+ cell line (black arrow). The sequestration of exogenous (fused with DsRed Ex fluorophore) or endogenous p53 (immune-labeled by TRITC) by exogenous HIF-1subunits (ECFP-HIF-1α and EYFP-HIF-1β) was observed in the form of nuclear foci (Fig. 4B). Moreover, sequestration of endogenous p53 by exogenous HIF-1 subunits in a concentration-dependent manner with subsequent enlargement in the size of foci further suggested integration of p53 and HIF-1 through direct physical interaction (Fig. 4C, Fig. S3A). Immuno-precipitation assays are difficult for three-component complexes and also, cannot be applied for the determination of homo-oligomeric constituents. Due to these reasons, we first improvised a variant of pull-down assay by replacing antibodies with biotin-labeled VEGF promoter to capture the complex upon chromatin and western blot steps with FRET by acceptor bleaching method to analyze interaction for the validation of p53, HIF-1α and HIF-1β as constituents of the complex. Next, we incorporated BN-PAGE based approaches to determine the homo-oligomeric state of p53. HCT116 p53−/− cells were co-transfected with the p53-DsRed Ex, ECFP-HIF-1α and EYFP-HIF-1β constructs and cell lysate was pulled by biotin-labeled promoter using streptavidin-coated agarose beads. Beads were washed to remove the non-specific binding and mounted on a glass slide for FRET detection by acceptor bleaching method (Fig. S3B). The bright spots on the beads were selected for bleaching and detection of FRET. The Fluorescence intensity of species on beads in −VEGF samples indicates insignificant non-specific pulldowns. The enhancement in ECFP and EYFP intensity in post bleach state of DsRed Ex confirms the interaction of p53-HIF-1 complex with VEGF promoter (Fig. S3C). Step-wise DsRed bleaching performed inside the cell showed similar enhancement in ECFP and EYFP intensity as observed upon agarose beads justifying pull-down of p53-HIF-1 complex by VEGF promoter (Fig. S3D). FRET on beads assay results is in agreement with the recent finding that p53 interacts with HIF-1 complex upon chromatin for adaptive responses in hypoxia (30). Next, the complex was checked against anti-p53, anti-HIF-1α, and anti-HIF-1β antibodies by two different modes in order to identify homo-oligomeric state of p53 that associates with the HIF-1. In the first mode, the same cell lysate was checked separately (Fig. 4D) or, the same immune band was checked in step-wise fashion (Fig. 4E). 3-12% and 4-16% gradients were used to resolve and detect other complexes with p53-HIF-1<M.W<p53-HIF-1. BN-PAGE immune blots clearly indicated cross immune reaction of the complex against three antibodies in hypoxic U20S p53+/+, HCT116 p53+/+ or MCF-7 p53+/+ samples confirming existence of p53, HIF-1α, and HIF-1β in the complex (blue arrows). Migration of p53 O and p53-HIF-1 in close proximity in 3-12% and 4-16% Bis-tris or 3-7% tris-acetate (Fig. S3E) gradient gel provides a possibility of p53 T as a constituent of the complex as p53T-HIF-1 has theoretical M.W. (≈424 KDa) comparable to p53 O. Therefore, in second mode, an innovative detergent displacement strategy was adopted with two-dimensional (2D) BN-PAGE method to confirm incorporation of p53 T. A combination of 18mM CHAPS, 0.1% Na-DOC and 0.1% TX-100 detergents (shown as D2) was determined in order to specifically dissociate constituents of p53-HIF-1complex. A separate anti-p53 or anti-HIF-1α BN-PAGE immune blot of the same hypoxic HCT116 p53+/+ cell sample showed the effect of D1 (18mM CHAPS), D2 or 0.5%TX-100 in cell lysis buffer over their respective higher-order assemblies. It is evident that D2 specifically dissociates p53 O into an intact T (magenta arrow). Besides, 0.5% TX-100 indicates the complete dissociation of p53T and HIF-1 from the complex (magenta or yellow arrow) and allowed higher-order HIF-1α species to enter into the gel (black arrows). However, p53-HIF-1 interaction seems to be partially intact in D2 with 0.1% TX-100 (blue arrows). Moreover, a diffused migration of p53 D was mostly observed with D2 (violet arrow, also see Fig. 6D) (Fig. 4F). In such a case, the approach to displace D2 combination in the cathode buffer during the second dimension run significantly dissociated the complex into p53T and HIF-1, which were identified based on their common spot in the merged anti-HIF-1α and anti-p53 2D BN-PAGE immune blots (Fig. 4G, yellow circles, also see Fig. 6D for merged p53 2D immune blot). Higher-order HIF-1α aggregates were visible in the second dimension too (black arrow).

**Figure4.**
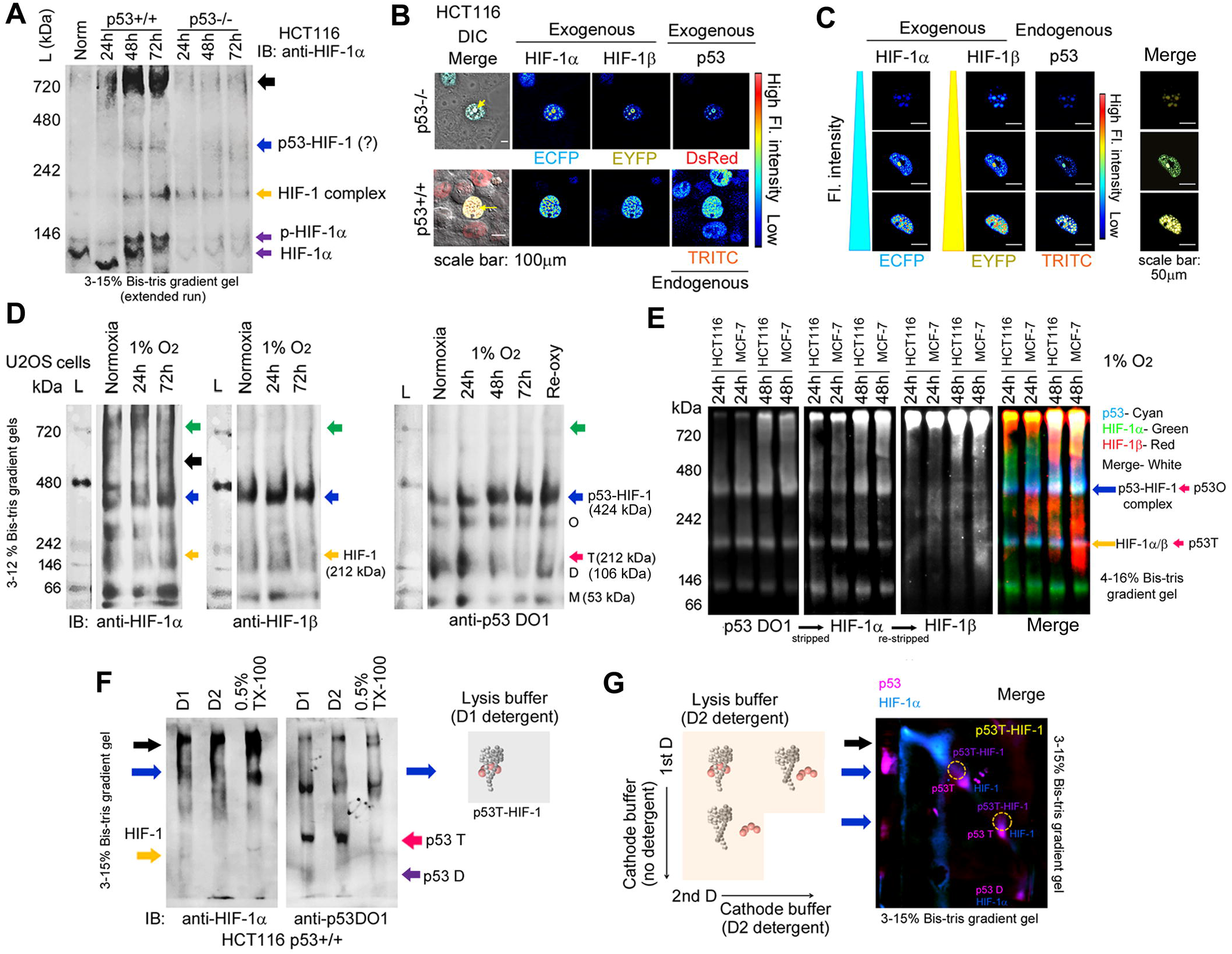
p53 T associates with HIF-1. **(A)** Anti-HIF-1α BN-PAGE immune-blot shows the rate of accumulation of different complexes of HIF-1α at 1% O_2_ in HCT116p53+/+ and HCT116p53−/− cells. Purple arrows indicate HIF-1α species (M.W. 120kDa), yellow arrow shows HIF-1 complex (M.W. 212 kDa) and blue arrow suggests p53-HIF-1 complex (M.W.>HIF-1) after an extended run of lysates in 3-15% Bis-tris gradient gel. The black arrow shows higher-order HIF-1α species in HCT116p53+/+ cell line. **(B)** Foci like structures (yellow arrows) showing co-localization of exogenous HIF-1α (ECFP), HIF-1β (EYFP) and exogenous or endogenous p53 (DsRed Ex or TRITC) in the nucleus of the cell. Scale bar 100μm. **(C)** Sequestration of endogenous p53 by exogenous HIF-1 subunits in concentration-dependent manner. Scale bar 50μm. Fluorescence images are pseudo-colored and color calibration bars indicate pixel-wise fluorescence intensity. **(D)** Triple immune reaction-based identification of endogenous p53T-HIF-1 complex. Green arrows indicate complex with M.W.>p53-HIF-1. The black arrow identifies higher order HIF-1α species. Blue, magenta and yellow arrows indicate p53-HIF-1, p53T and HIF-1 complex respectively. Native protein standards were separated from the PVDF membrane post-transfer and stained separately by Coomassie G250. **(E)** Identification of endogenous p53-HIF-1 complex by cross-reaction of the same immune band against three antibodies by stepwise stripping. anti-p53 DO1 (cyan), anti-HIF-1α (green) and anti-HIF-1β (red) immune blots were merged cautiously *in silico* to detect cross-reactivity (white). **(F)** Effect of different detergent combinations on p53 or HIF-1α complexes. Blue arrows indicate p53-HIF-1 complex positions in the immune-blots. Anti-p53 immune-staining confirms dissociation of intact T from p53-HIF-1 complex by D2 detergent (magenta arrow). **(G)** Schematic representation of the principle of detergent displacement strategy (left panel). Anti-HIF-1α immune blot was stripped for anti-p53 immune detection and two immune blots were cautiously merged *in silico* to identify the dissociated p53T (magenta) and HIF-1(cyan) entities (dotted yellow circles) (right panel). Higher-order HIF-1α aggregates are shown by black arrows. For the merged anti-p53 immune-blot image, refer to Fig 6D. 3-15% Bis-Tris gradient gel was selected for proper resolution of all complexes in 1D and 2D BN-PAGE run.

**Figure5.**
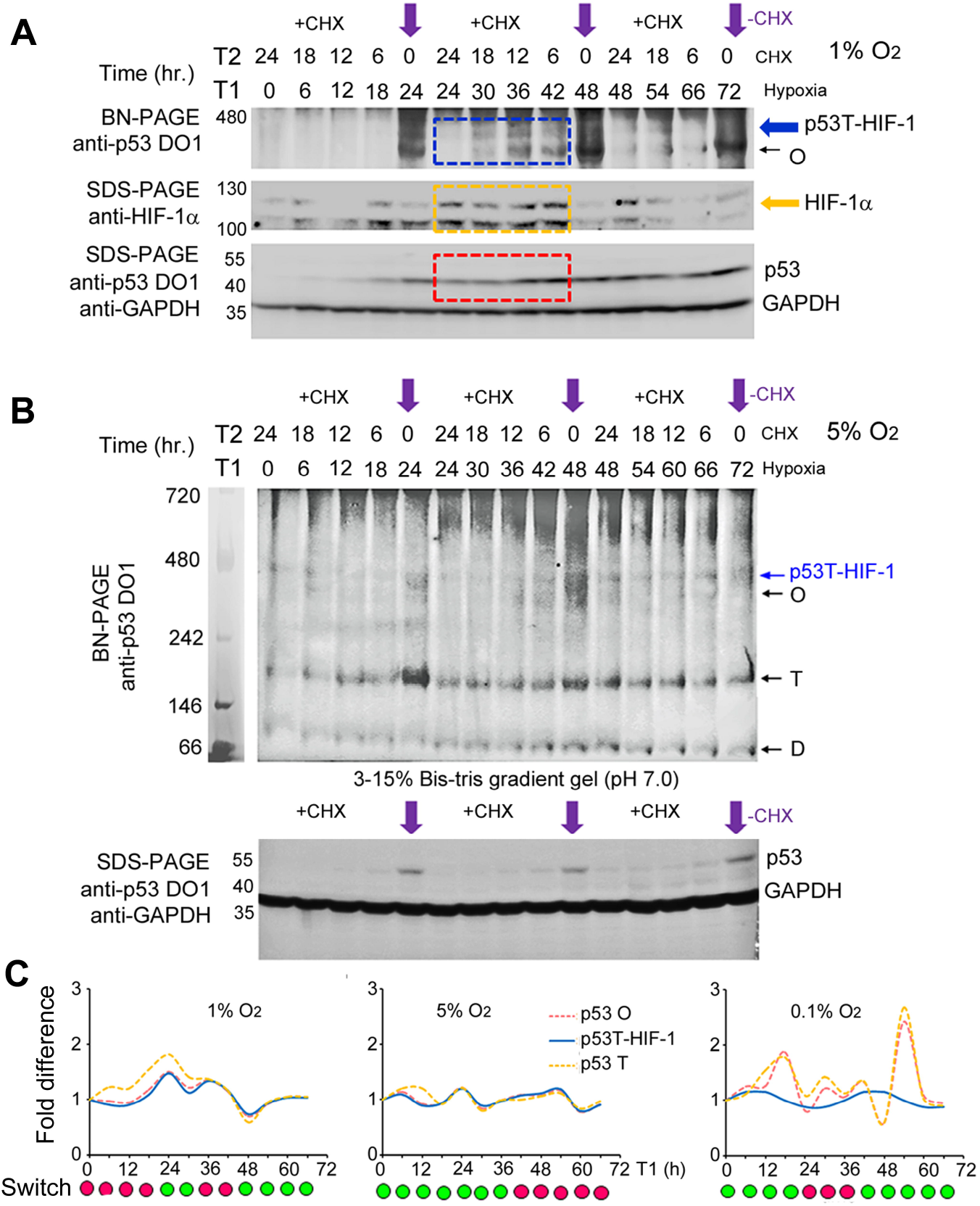
*off* state of T⇀O switch controls p53T-HIF-1 complex generation. **(A, B)** p53T-HIF-1 interaction was analyzed by the comparison of anti-p53 and anti-HIF-1α SDS-with anti-p53 BN-PAGE immune-blots at 1 (A), 5 (B) or 0.1% O_2_ (Fig S3F). Blue, yellow and red boxes indicate p53T-HIF-1 interaction. **(C)** Fold change in BN-PAGE p53T-HIF-1 complex signals at each interval indicates that T and HIF-1 association is favored in *off* mode of the switch. The rate of the p53T-HIF-1 complex association was measured from the anti-p53 BN-PAGE immune blots (rather than anti-HIF-1α blots) for the quantitative comparison with p53 T and O. At 0.1% O_2_, p53 T & O are favored as compared to p53T-HIF-1 complex. Due to the inclusion of protein standards in 15 well gel, 60^th^ h sample for 1% O_2_ was analyzed separately or from other replicates. Native protein standard shown for 5% immune blot was run simultaneously in a different gel of the same gradient.

**Figure6.**
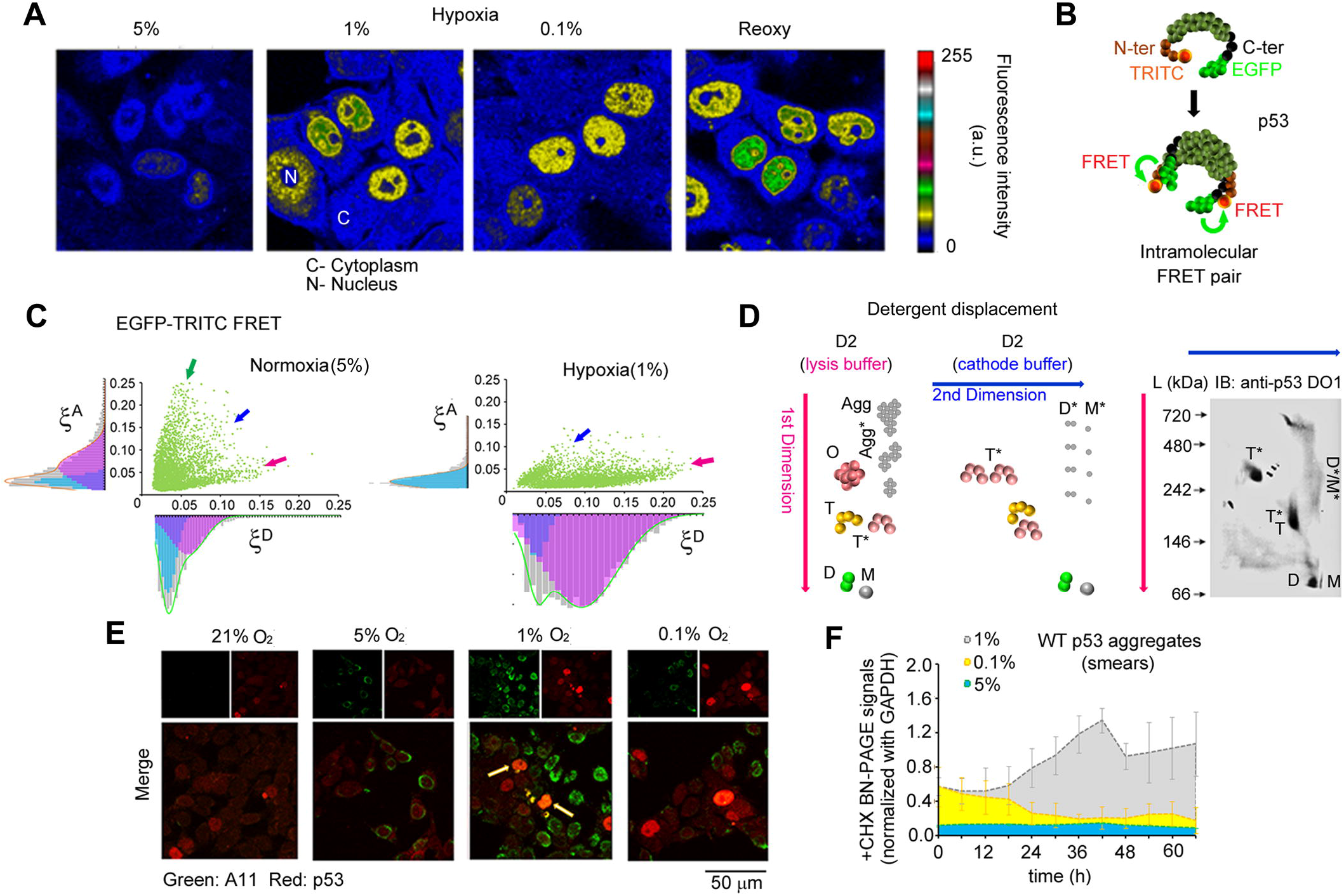
Conformation change in D is linked to prion-like aggregation of WT p53. **(A)** Sub-cellular localization of p53 in the hypoxia gradient. IFC staining shows localization of p53 in the cytoplasm(C) and nucleus (N) in hypoxia gradient. The color calibration bar represents pixel-wise fluorescence intensity. **(B)** F-TEPA for the determination of conformation change associated with p53 homo-oligomers. **(C) ξ** ^D^ v/s **ξ** ^A^ scatter plot indicates to the conformation change in lower-order p53 homo-oligomer D or M (magenta arrow) in hypoxia. T (blue arrow) or O (green arrow) remains conformationally intact in both normoxia and hypoxia. For **ξ**^D^ & **ξ**^A^ histograms and scatter plot generation, all the calculated **ξ**^D^ & **ξ**^A^ images (n≤20) from three independent replicates were used to randomly select at least 5000 pixels by automated methods. **(D)** 2D BN-PAGE by detergent displacement strategy shows the disintegration of p53 aggregates into M*/D*. anti-A11 staining (immune-labeled by FITC) of endogenous p53 (immune-labeled by TRITC) in HCT116 p53+/+ cells under variable oxygen. Arrows in merged images show p53 amyloid aggregates detected by A11 antibody. **(F)** CHX trap based BN-PAGE analysis of p53 aggregates (observed as smears) under variable oxygen (for immune-blot refers to Fig. 2B, C or Fig 5B). The value represents mean and error bar shows s.d. from three independent experiments.

To confirm whether p53T-HIF-1 is the result of the function of T⇀O switch, its dynamics was monitored in HCT116 p53+/+ cells by CHX-trap based BN- or SDS-PAGE methods in hypoxia gradient. Fold difference in the level of p53 T, O or p53T-HIF-1 complexes at each interval indicated that the o*ff* state is preferred for p53T-HIF-1 generation at 1 or 5% O_2_. p53T-HIF-1 was abrogated under sustained *on* state at 0.1% O2 (Fig 5A-C, Fig S3F, G) confirming that p53T-HIF-1 dynamics is directly controlled by T⇀O switch.

### In off mode, a reversible tetramer⇌ dimer state controls p53 prionoid and chaperone function

Enhanced cytoplasmic localization of p53 (Fig. 6A and Fig S4A) in addition to the appearance of smears (D<M.W.<T/O) (see Fig. 2B, red arrow) directs towards the possibility of prionoid function of p53 (31) via hypoxia-induced conformation change (3) in the *off* state of the switch. Besides, the enhanced p53T-HIF-1 association provides clue to the chaperoning of flexible DNA-reading head of HIF-1(32) by p53T (see Fig. 5C). In order to prove that a reversible D⇌T system controls two antagonistic functions of p53 via **D**⇌**T**↽O shift in *off* mode, first, we employed an improvised FRET-based terminal ends proximity analysis (F-TEPA) to identify if p53 functions like a prion upon conformation change in hypoxia. FRET variation between terminal ends (N- and C-ter) of p53 quantified in terms of two quantities **ξ**^D^ and **ξ**^A^ (See materials and methods) in F-TEPA provided information regarding conformation change associated with p53 homo-oligomers in hypoxia (1% O_2_) (Fig. 6B). p53-EGFP construct (fused at C-ter) was transiently transfected in HCT116p53−/− cells and subjected either to normoxia (5% O_2_) or 24h hypoxia (1% O_2_). The conformation of p53 molecules was captured by chemical fixation of cells post-exposure and N-ter was indirectly immune-labeled with primary anti-p53 DO1 (N-ter specific monoclonal antibody) and TRITC tagged secondary antibodies. The fluorescence intensity of EGFP, TRITC, and FRET was acquired by spectral imaging and linear unmixing in order to calculate the pixel-wise distribution of **ξ**^**D**^ & **ξ**^**A**^ proximity coefficients. **ξ**^**D**^ represents FRET efficiency due to donor quenching that corresponds to conformation change in p53 molecules. **ξ**^**A**^ signifies fraction of acceptor sensitized in particular assemblies and thus relates to p53 homo-oligomerization states.

The pixel-wise histograms of **ξ^D^**& **ξ^A^**(fitted with a sum Gaussian components) from EGFP-TRITC FRET system were aligned to their respective axes in **ξ**^**D**^ v/s **ξ**^**A**^ scatter plot. The associated scatter plot relates to the conformation change in p53 homo-oligomers (Fig. 6C). As CHX trap based BN-PAGE immune blot showed stabilization of p53 mostly in D with low amount of T at 1% (see Fig. 2B, compare D and T accumulation in lane 6) and exist in mixed state of D, T and O forms at 5% O_2_ or normal state, therefore, slopes appeared in **ξ**^**D**^ v/s **ξ**^**A**^ histogram by F-TEPA could be linked to the conformation of D (magenta arrow), T (blue arrow) and O (green arrow) in normoxia and hypoxia. The decrease in **ξ**^A^ with an increase in **ξ**^**D**^ indicates conformation change in lower-order p53 homo-oligomers (D/M) under hypoxia when compared to normoxia. Higher-order p53 assemblies remained dominantly in the native state. 2D BN-PAGE by D2 detergent displacement strategy showed disintegration of p53 aggregating smear (see Fig. 2B, red arrow) into M*/D* by anti-p53 immune-blotting. Since D2 does not affect T and D^ML^ is the most dominant form in the hypoxic pool at 24h, therefore analysis proved that D^ML^ triggers p53 self-aggregation and thus confirmed prionoid function of p53 (Fig. 6D). Anti-A11 IFC staining in combination with densitometry of aggregating smears by CHX-trap based BN-PAGE further identified that amyloid aggregation induced by D^ML^ is preferred in *off* mode of T⇀O switch (Fig. 6E, F).

**ξ**^D^ and **ξ**^A^ histograms showed conformational dynamics of D^ML^ by FITC-p53-DsRed Ex F-TEPA system at different intervals in hypoxia (Fig. 7A). It is clear from the comparison of **ξ**^D^ (Fig. 7B) and **ξ**^A^ (Fig. 7C) components that D^WT^⇀**D**^**ML**^ (in the initial 0-24h interval) reverses to **D^WT^**↽D^ML^, similar to re-oxygenation, and triggers D⇀T in later duration (24-72h). At 72h, an equilibrium D^**WT**^⇌T state is achieved as observed through histograms. When compared to R.A. measurements by CHX-trap based BN-PAGE in hypoxia (compare with Fig. 2D, middle panel), it is clear that **D^ML^**↽D^WT^**↽**T state controls prionoid function, and a reverse D^ML^⇀D^WT^**⇀T** controls p53T-HIF-1 complex via a reversible D⇌T state in *off* mode.

**Figure7.**
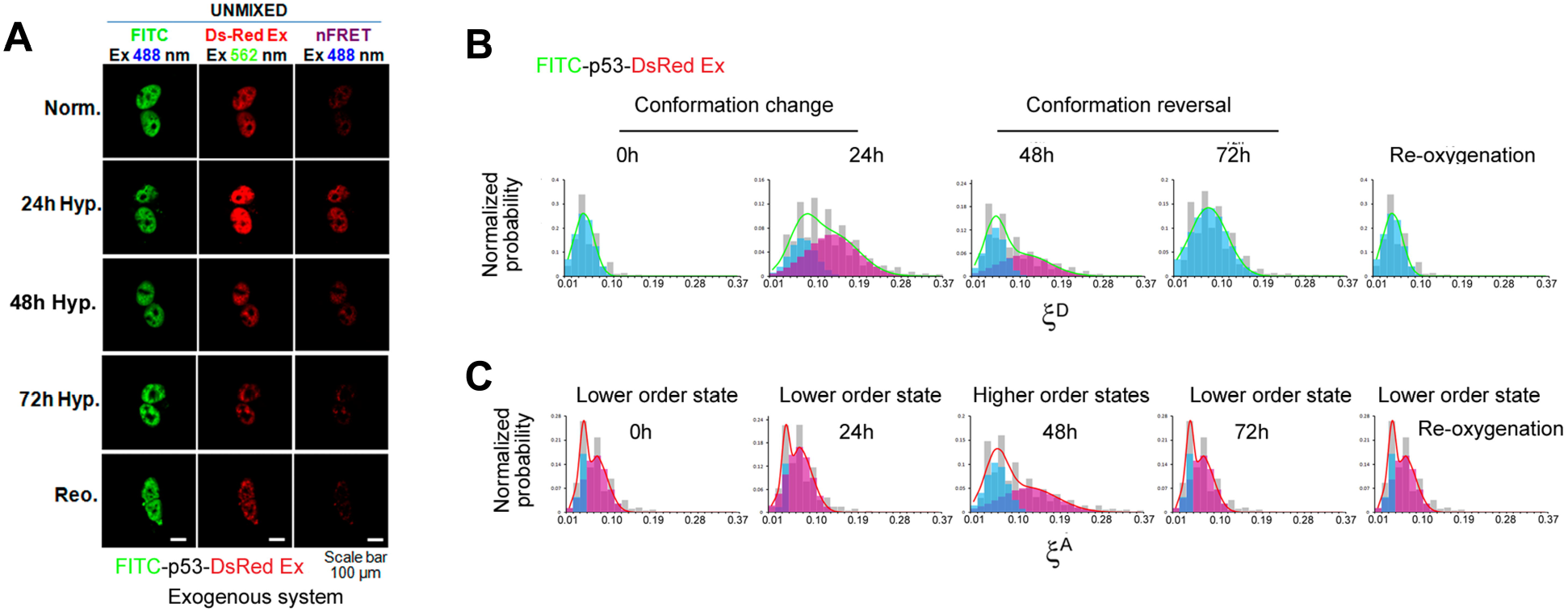
Conformational dynamics of mutant-like p53 D in hypoxia. **(A)** Time-dependent F-TEPA analysis performed in hypoxia (1% O_2_) using FITC and DsRed Ex FRET pair generated at N- and C-ter of exogenously expressed p53 in HCT116 p53−/− cells. Scale bar 100μm. **(B, C)** The pixel-wise **ξ**^D^ and **ξ**^A^ histograms for FITC-p53-DsRed Ex F-TEPA system show conformation change (depicted by higher **ξ**^D^) of p53 D (depicted by lower **ξ**^A^) after 24h hypoxia exposure. In the 24-72h interval, a decrease in **ξ**^D^ and enhancement in **ξ**^A^ indicates conformation reversal of D^ML^ species and the formation of T (compare with Fig. 1D, middle panel). The histogram represents measurements from 5000 pixels selected randomly from n≤20 **ξ**^D^ & **ξ**^A^ images (512×512 pixel^2^) of cells from all the independent experiments (n≥3). Re-oxygenation was used as positive control.

Crystallographic studies have shown that N-ter of HIF-1α and HIF-1β exists in a constricted conformation which extends upon DNA for optimal binding of HIF-1 complex (32). To check whether p53 assists as a chaperone for this process in the cell, first, **ξ**^D^ histogram from the F-TEPA system constituting ECFP-HIF-1α and EYFP-HIF-1β (both fluorophores are fused at N-ter) was compared in HCT116 p53+/+ and HCT116 p53−/− cells. As p53, HIF-1α and HIF-1β are co-localized in nuclear foci; results indicated stabilization of HIF-1 complex by p53 in the distal configuration of N-ter (DNA-reading head) of its subunits irrespective of hypoxia (Fig. 8A-C). An introduction of DAPI as FRET donor in F-TEPA systems was used to determine whether stabilized HIF-1 efficiently binds to DNA upon conformational modulation by p53. DNA-HIF-1 interaction was analyzed by DAPI-EYFP FRET pair in HCT116 p53+/+ and HCT116 p53−/− cells (Fig. 8D). ECFP-HIF-1α was replaced by non-fluorescent HIF-1α in both hypoxic HCT116 p53+/+ and HCT116 p53−/− cells to eliminate ECFP contamination due to cross-excitation by 403nm (used to excite DAPI). EYFP foci represent HIF-1 complex as both subunits colocalize in it (Fig. 8E). Spectral imaging and linear unmixing were performed to determine FRET-based detection of HIF-1-DNA association through pixel-wise **ξ**^A^ distribution histogram analysis. HCT116 p53+/+ indicates higher DAPI-EYFP **ξ^A^** as compared to HCT116 p53−/− cells confirming p53-dependent stabilization of HIF-1 upon DNA (Fig. 8F). ECFP-HIF-1α was utilized along with DAPI and EYFP-HIF-1β in HCT116 p53+/+ cells in order to analyze oxygen-dependent HIF-1 conformational modulation by p53 upon DNA (Fig. 8G). Due to simultaneous excitation by 403nm laser, DAPI & ECFP combination can be regarded as one donor for an acceptor EYFP in order to analyze HIF-1-DNA interaction via (DAPI+ECFP)-EYFP FRET pair. To observe the change in EYFP sensitization, spectral imaging and linear unmixing were performed. ECFP* indicate unmixed ECFP emission in (DAPI+ECFP) combination (Fig. 8H). For such a pair, it was easier to interpret results via **ξ^A^** histogram as it excluded complexity due to contributions from two simultaneously excitable donors. Histograms revealed a decrease in **ξ^A^** under hypoxia when compared to normoxia (Fig. 8I). When compared to Fig. 8C, it suggests that p53 further brings alteration in distal HIF-1 subunits upon DNA to enhance efficiency of HIF-1-DNA interaction (see Fig. 8F) in response to fluctuating oxygen.

**Figure8.**
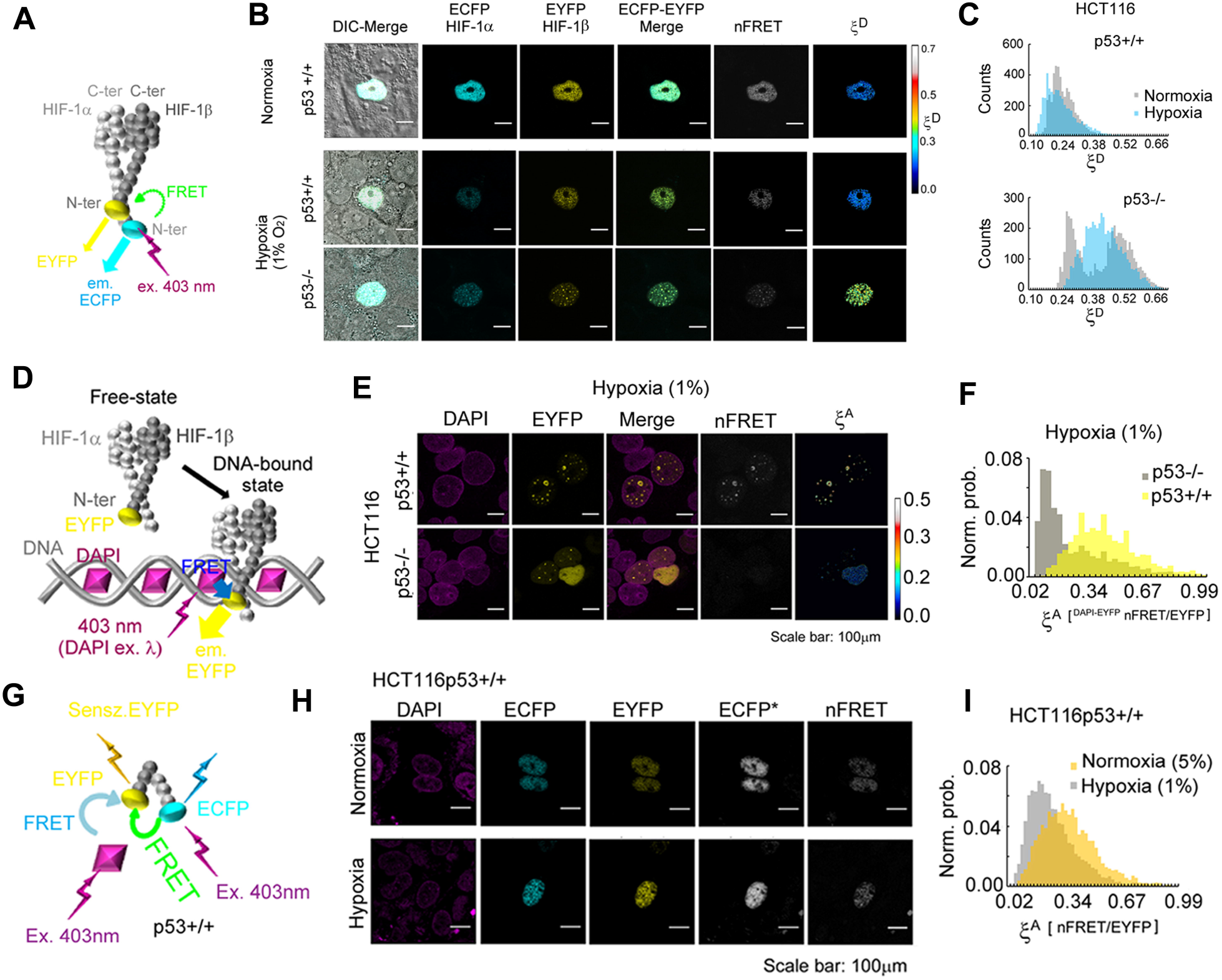
Modulation of HIF-1 complex by p53. **(A)** F-TEPA analysis with exogenously co-expressed ECFP-HIF-1α and EYFP-HIF-1β. **(B)** Confocal images showing ECFP, EYFP, nFRET and **ξ**^D^. Images were obtained by spectral imaging and linear unmixing. Color calibration bar indicates pixel-wise **ξ**^D^ values. Scale bar 100μm. **(C) ξ**^D^ histogram shows HIF-1 stabilization by p53 irrespective of hypoxia. Lower **ξ**^D^ for HCT116 p53+/+ cell line indicates distal HIF-1 subunits in the presence of p53. The histogram represents measurements from 5000 pixels selected randomly from n≤20 **ξ**^D^ images (512×512 pixel^2^) of cells from all the independent experiments (n≥3). **(D-F)** p53 dependent stabilization of HIF-1 complex upon chromatin in hypoxia was determined by the inclusion of DAPI as a donor in the FRET system. Non-fluorescent HIF-1α was utilized with fluorescent EYFP-HIF-1β subunit in isogenous HCT116 cell lines and DAPI-EYFP FRET was quantified in terms of **ξ**^A^. Scale bar 100μm. **(G-I)** Oxygen-dependent modulation in conformation of HIF-1 complex by p53 was determined by the incorporation of ECFP-HIF-1 in the FRET system used in (D-F). ECFP* is the pure ECFP emission after simultaneous excitation of (DAPI+ECFP) donor combination. Scale bar 100μm. The histogram in (F,I) represents measurements from 5000 pixels selected randomly from n≤20 **ξ**^A^ images (512×512 pixel^2^) of cells from all the independent experiments (n≥3).

In order to verify p53-dependent conformation change in HIF-1, a three chromophore F-TEPA system was generated in HCT116 p53+/+ cells with fluorescent HIF-1 subunits and TRITC labeled N- or C-ter of endogenous p53. Scatter plots represent pixel-wise **ξ**^D^ values for FRET between HIF-1 subunits and p53 C-ter (upper panel) or p53 N-ter (lower panel) in normoxia (5% O_2_) and hypoxia (1% O_2_). It is evident that p53 C- and N-ter are close to HIF-1α and HIF-1β in HIF-1 complex respectively (as shown by a dotted ellipse in the two scatter plots). However, upon transition to hypoxia, separation of p53 N-ter and HIF-1β is specifically decreased (as indicated by an increase in **ξ**^D^, blue arrow) (Fig. 9A) indicating stabilization of a flexible HIF-1 subunit (32). Data show p53 as oxygen-dependent chaperone which further adjusts separation of distal HIF-1 subunits to modulate HIF-1-DNA interaction under variable oxygen-crisis. Luciferase assays indicated that such an arrangement activates HIF-1 upon HRE (hypoxia response element) (Fig. S4B). On the contrary, comparison of p21(Fig. S4C, D) and p53T-HIF-1 dynamics at 5% O_2_ (see Fig. 5C, middle panel) suggested that a more proximal arrangement of HIF-1 subunits would allow p53 self-activation via T-DNA interaction under normoxic conditions. Collectively, data suggest that p53T may self-align and chaperone HIF-1 upon DNA in order to segregate self- from HIF-1 activation depending upon oxygen crisis in *off* mode of the switch (Fig. 9B). It would be tempting now to decipher the role of the p53T-HIF-1 complex upon DNA damage stress in inducing p53-mediated cell fate decisions. In conclusion, *off* mode controls both prionoid and chaperone function of p53 via reversible D⇌T step. Taken together, our finding invariably proves p53 as oxygen-sensor that operates via T⇀O switch in segregating its role of a tumor suppressor or promoter in tumorigenesis (Fig. 9C).

**Figure9.**
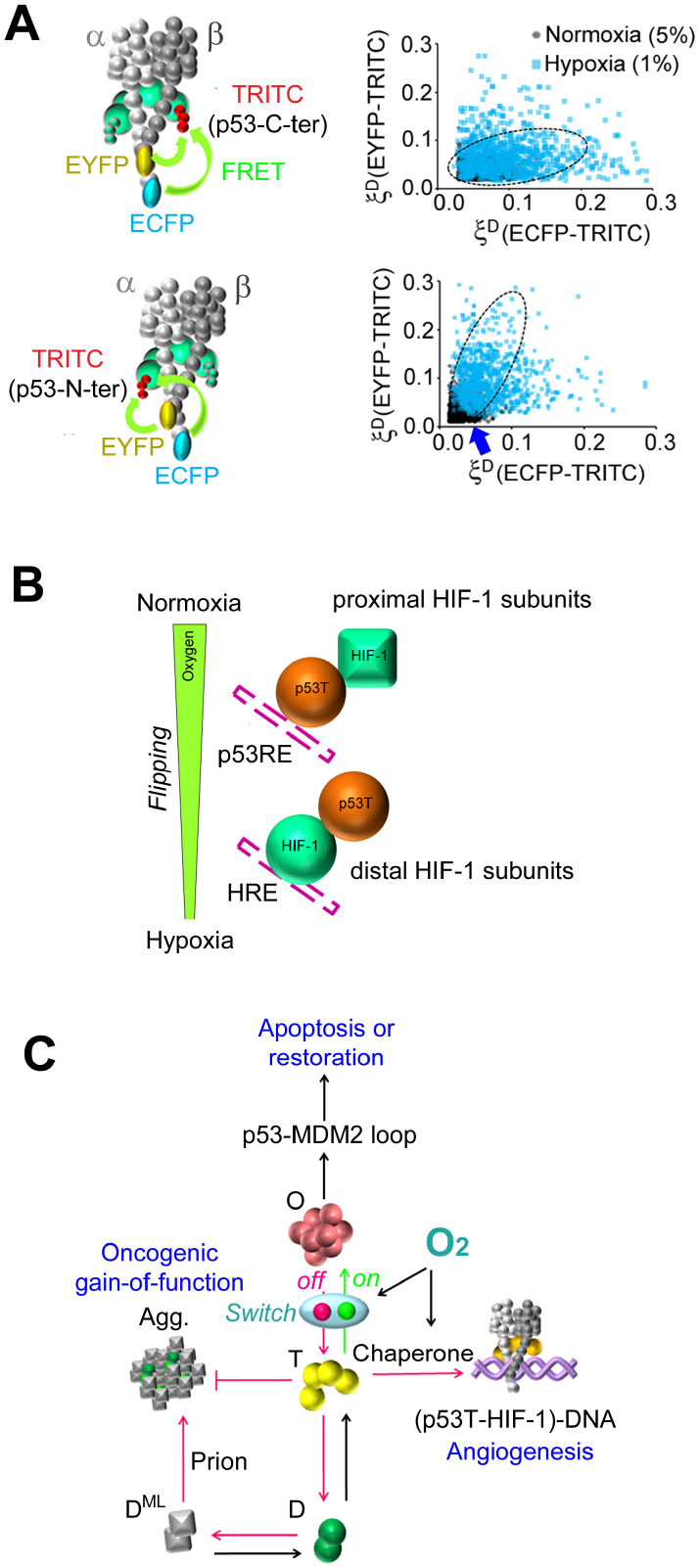
p53 chaperones HIF-1 complex. **(A)** Three chromophores F-TEPA shows chaperoning of HIF-1 complex by p53 in an oxygen-dependent manner. **(B)** A possible Flip-flop mechanism by which p53T flips upon self and HIF-1 response elements (REs) by virtue of its chaperone function. **(C)** Model showing operation of oxygen- responsive p53 T⇀O switch in dictating cell fate. The model does not strictly represent the structure of protein complexes.

## DISCUSSION

Low oxygen disturbs homeostasis and usually manifests in multiple pathophysiological conditions including cancer. p53 is linked to hypoxic tumor progression through its inactivation and loss of apoptotic potential in the mutant-like state (2). However, as an oxygen-sensor, how p53 transforms its inactive state, switches the function of a tumor suppressor and segregates its prionoid and chaperone functions, remains questions of prime importance under variable oxygen crisis.

Through an innovative “homo-oligomer trap” design by CHX and MG132 based interventions during normal proliferation of cells, we have identified a metastable state of p53 T by BN-PAGE which justifies a possibility of D↽T and T⇀O as functional switches upon stimulation. The dissociation of T into D exclusively after inhibition of complete cellular protein synthesis suggests that D↽T switch could operate by the involvement of other entities that would drive D-D association via post-translational modifications. On the contrary, the association of T into O specifically under the diminishing protein degradation in cells, a condition similar to hypoxia, indicates that T⇀O switch could be oxygen-sensitive and its independency of protein synthesis rate suggests that such operation may not involve the same modifications as D⇀T. Through R.A. measurements, we show that T⇀O switch operates upon D⇌T⇌O and transforms it into D⇀T⇌O and D⇌T↽O states in its *on* and *off* modes respectively thus calibrating stabilized p53 pool in a gradient of hypoxia. Duration of the switch seems crucial in determining the strength of these two homo-oligomerization states that may ultimately switch the role of p53 either as a tumor suppressor or promoter in dictating cell fate.

It’s well known that D⇀T rate controls the p53-MDM2 feedback loop for cyclic or terminal cell fate (13, 20, 26). Therefore, an *on* mode D⇀T⇌O shift could be responsible for p53’s tumor-suppressive function via D⇀T step. However, being linked to p53-MDM2 loop activation, we discovered that a stable T⇌O state is essential for this role in an oxygen-dependent manner. p53 O state is well characterized to consist of two tetramer units and known to interact with DNA (18). In addition, p63 or p73 has also been shown to exist in octamer assemblies (19). Our detergent displacement strategy also showed dissociation of O into T by 2D BN-PAGE. However, their functions are less understood. p53 O is found to be ubiquitinated in the basal pool and therefore, is more likely to induce MDM2-mediated p53 restoration and maintain basal state via T⇀O step. On the contrary, a stable O may activate p53 for its target gene expression, including MDM2 resulting in apoptosis via T↽O step. Restoration and activation cycles are crucial for p53 dynamics and T⇌O might play an important role in shaping p53 digital mode into sustained or repetitive pulse for a flexible cell fate decision upon stress (1). Further, p53 is also known to induce a variable cell fate response due to differential ubiquitination (33). In such a case, O ubiquitination might play an important role in promoting apoptosis or growth arrest in *on* mode. Besides, such a phenomenon might also control O and DNA interaction for expanding the repertoire of p53 REs (response elements) (18) and modulate T-DNA association for distinct outcomes via differential expression of p53 target products.

By adopting a novel “detergent displacement” strategy with 2D BN-PAGE and performing innovative F-TEPA with multiple fluorophores, we show that an *off* mode D⇌T↽O shift can segregate p53 characters of a prion and chaperone. Under D↽T, reversible transitions through D^WT^⇌D^ML^ conformational states, the mechanism for which remains still unknown, can initiate wild-type p53 aggregation and oncogenic gain-of-function (19) in a prion-like fashion. On the other hand, D⇀T can promote T-HIF-1 complex which displays a chaperone function of p53. Recently, it has been shown that WT and MT p53 binds to HIF-1 complex at chromatin for its stabilization upon HRE in controlling angiogenesis and other HIF-1α induced transformations in hypoxic tumor progression (30). Our three chromophore FRET setup identifies a distal and proximal arrangement of two HIF-1 subunits, α and β in hypoxia and normoxia by the presence of p53 respectively. A simple introduction of DAPI in three chromophore setup further identified that a distal configuration of two subunits is suited for HIF-1 binding whereas a proximal configuration is favorable for p53T binding upon chromatin. Hence, a “flip-flop mechanism” could be hypothesized by which T flips upon HRE and p53RE by virtue of its oxygen-sensitive chaperone character and modulates self or HIF-1 activation via a common T-HIF-1 complex in a hypoxia gradient. Such a mechanism may be beneficial in balancing expressed gene products of both entities through their common transcriptional co-activator p300 (34) and regulating tumorigenesis in normoxia and hypoxia.

It seems that such a transcriptional control is dependent upon p53T-HIF-1 stabilization in synchronous with p53T in response to stress. In hypoxia (1% O_2_), the sustained pattern can induce HIF-1 target genes (e.g. VEGF) to initiate hypoxia adaptive processes via p53T-(HIF-1-DNA) association. Conversely, in normoxia (5% O_2_), a transient profile can manage basal levels of p53 target genes (e.g. p21) via flipped (DNA-p53T)-HIF-1 complex. p53’s spontaneous pulse is transcriptionally attenuated that does not allow p53 to transcribe p21 above base level and interfere with the progression of cells (35), in which case p53T-HIF-1 dynamics could be responsible for the maintenance of baseline and lowering of p21 threshold when abrupt fluctuations in p53 are encountered in normal growth conditions. A detailed investigation at single-cell level with smaller time intervals may answer whether transition from transient to sustained stabilization pattern of p53T-HIF-1 leads to p53’s product expression above basal level (threshold), in a manner similar to well-characterized p53 dynamics (1). Fold difference in CHX-trap based BN-PAGE immune density level at each interval has shown overlapping p53T and p53T-HIF-1 dynamics at 1 and 5% O_2_ although the trend is disrupted at 0.1% O_2_. This observation suggests that hypoxia gradient can modulate the *on-off* duration of T⇀O switch in order to synchronize p53 and p53T-HIF-1 for synergistic or antagonistic interplay of p53 and HIF-1α in tumorigenesis.

The reversible T⇌O and D⇌T states may generate auto-inhibitory effects. In *on* mode, p53 restoration via T⇀O can inhibit p53 activation via T↽O during D⇀T⇌O shift. Similarly, in *off* mode, p53 prion effect via D↽T can restrict p53 chaperone function via D⇀T during D⇌T↽O shift. Therefore, to generate a stable response, control over D⇀T and T↽O conversions are crucial as they can regulate the amount of metastable T that may perturb T⇌O and D⇌T equilibrium states and limit auto-inhibitory effects in order to ultimately switch p53 function of a tumor suppressor or promoter in hypoxia gradient. It may be possible that oxygen-dependent post-translational modifications are involved in the fine-tuning of D⇀T and T↽O regulations (36).

Although genetic p53 mutants have been earlier shown to drive more than 50% of all cancer that can initiate prion-like aggregation of self (31) or other tumor suppressors like p63 and p73 in decreasing life expectancy of patients (19), it is unknown why a higher percentage of them also carry on wild-type p53 allele. Perhaps, p53 heterogeneous alleles have the inbuilt information in order to *on* and *off* the process of decision making under stimuli. Under stress, the T⇀O switch, in a generalized fashion, could control the cell’s orientation program by transforming metastable p53 homo-oligomerization states at ease. It is then important to ask if, during the early stages of tumorigenesis, the T⇀O switch might be operational in triggering cell competition (37) between oxygenated and oxygen-deprived cells. Our discovery that p53 behaves like a cellular oxygen sensor via T⇀O switch has implications beyond cancer and futuristic intervention via oxygen manipulation may reverse cell’s aberrant behavior in hypoxic tumors and other lifestyle diseases.

## EXPERIMENTAL PROCEDURES

### Cell culture and Hypoxia treatment

Human colon cancer HCT116 p53+/+, HCT116 p53−/−(38,39), U2OS p53+/+ cells (NCCS, Pune), MCF-7 p53+/+ (NCCS, Pune) were cultured in Dulbecco’s modified Eagle’s medium supplemented with 10% (v/v) FCS, penicillin (100U/ml), streptomycin (100U/ml) and L-glutamine (4mM), pH 7.4. For the proliferating cell cultures, 5% CO_2_ was maintained in a humidified incubator at 37 °C. Hypoxia exposure to the cell culture was performed under controlled conditions in a hypoxia chamber (Plas Labs inc., USA) equipped with a thermostat, O_2_ and CO_2_ sensors. O_2_ and CO_2_ levels were quickly restored upon the detection of fluctuations (by ±0.2units) from the set levels through the automated purging of gases by inbuilt sensors. The normal atmospheric pressure was maintained through the N_2_ gas. Just before the exposure, DMEM was replenished in the cultures to sustain 72h exposure without significant change in pH (7.2-7.4). Re-oxygenation was performed by transferring the cultures from the hypoxia chamber to the CO_2_ incubator for a 24h period.

### Generation of CHX-trap (Homo-oligomer trap) in hypoxia

p53 appears in spontaneous oscillation cycles during normal growth of the cell. In a cell population, these cycles are asynchronous and when captured by population methods such as BN- or SDS-PAGE, usually average out the actual fluctuations in p53 dynamics. Besides, a minimal pool generated by asynchronous cycles at a given time can further hinder the detection of p53 dynamics. Complete inhibition of total protein synthesis and degradation in the cell population has an advantage in suppressing average out effect. The average out effect arises from heterogeneous distribution of p53 pulse in its initiation, stabilization or degradation phase and their interference at a single-cell stage in a population. In principle, inhibition of degradation can amplify p53 pool from initiating as well as degrading pulses. On the contrary, protein synthesis inhibition can provide major contributions from stabilized pulses as it inhibits initiation and allows degradation. Together, these interventions can result in amplification of p53 pool with a degradable (now stabilized) and stabilized pulse suppressing negative interferences, and hence average out effects in a population. BN-PAGE performed after such manipulation provided p53 homo-oligomerization in its stabilized state with improved detection under a given condition that successfully identified the ubiquitination of octamer during p53 restoration in basal state or different equilibrium shifts in hypoxia gradient via the switch.

For the generation of CHX-trap in hypoxia, HCT116 p53+/+ cells were seeded in three groups of four six-well culture dishes 24h before the assay. For the assay in 1% O_2_, plates of the 1^st^ group (with approx. 5.0×10^5^cells in each well) were sequentially exposed to the hypoxia at 6h intervals followed by the plates of the 2^nd^ (2.0×10^5^cells) and 3^rd^ groups (3.0×10^4^cells). After the last plate of the 2^nd^group was exposed, Cycloheximide (100μM) treatment was initiated from the first to last exposed plates of all the three groups at 6h intervals. The experiment was designed for the simultaneous processing of samples at the end of the hypoxia treatment with the maintenance of the 24h≥ CHX treatment ≥6h duration at each interval as well as the capture of the protein dynamics from 0-72h at 6h intervals. For the assay under 5% and 0.1% O_2_, only the number of cells in the initial seeding process differed considering the normal proliferation and apoptosis effects under these conditions. Cells in each well of the plate were pooled for further analysis by SDS- or BN-PAGE immune blotting after the exposure.

To capture the dynamics of p53 homo-oligomers under normal growth conditions, complete cellular protein synthesis and degradation were inhibited by CHX and MG132 respectively in order to trap spontaneous p53 oscillations. However, in hypoxic conditions, MG132 was excluded as hypoxia itself perturbs p53 degradation.

### BN- and SDS-PAGE

Following the hypoxia treatments, cells were rinsed with 1X TBS (20mM Tris-HCl, 150mM NaCl, pH 7.4) inside the hypoxia chamber. 1X TBS with DNase, RNase, protease and phosphatase inhibitors at room temperature (RT) were used as common compositions for the lysis buffer in each experiment. 18mM 3-[(3-cholamidopropyl) dimethylammonio]-1- propane sulfonic acid (CHAPS) was used as a detergent (as D1 in the main text) in the base buffer in all BN-PAGE experiments (19) except for the 2D BN-PAGE. Here, 18mM CHAPS was supplemented with 0.1% TritonX-100 (TX-100) and 0.1% sodium deoxycholate (Na-DOC) as detergents (as D2 in main text). For each 6 well plates, 50μL lysis buffer was added in each well and cells were pooled in the single micro-centrifuge tube and transferred to the desk. The lysis was allowed for 20 min. followed by centrifugation and collection of the supernatant at RT. For 1D BN-PAGE experiments, the supernatant was added with 20% glycerol and 5mM Coomassie G-250. For SDS-PAGE experiments, 2% SDS was further added and samples were denatured at 95°C for 10 min with mercaptoethanol. The samples were estimated for protein concentration by Bradford protein assay kit (BioRad) and loaded on to the gradient gels in equal masses. Gel buffer with 50mM Bis-Tris, 50mM ε-aminocaproic acid, pH 7.0 was used with both lower and higher acrylamide-bisacrylamide (32:1 with 40% acrylamide) mix for the casting of Bis-Tris (pH 7.0) or Tris-acetate (pH 7.0) native gel gradient. For 1D BN-PAGE, electrophoresis of the samples was performed in cathode buffer containing 15mM Bistris, 50mM Tricine, and 0.02% Coomassie blue G250 and anode buffer with 50mM Bistris (pH 7.0) at RT. NativeMark protein standards were run with the samples for the identification of molecular complexes based on the mol. wt. In 2D BN-PAGE, a lane of 1D BN-PAGE was cut from the gel and mounted horizontally to the top of 3-15% Bis-Tris gradient gel. The complexes were then segregated with Na-DOC(0.1%) and TX-100(optimizable in the range of 0.001-0.1%) in cathode buffer by a 2D run at RT. All the BN-PAGE gels were run at constant voltage (60V). For SDS-PAGE, samples were fractionated by hand made 10% Bis-Tris gels in MOPS-SDS running buffer containing 0.1% SDS, 50mM MOPS and 50mM Tris-Base, pH 7.0 at constant 120V at RT. After the SDS- or BN-PAGE based fractionation, polyacrylamide gels were soaked in 10% SDS for 30 min at RT. The proteins were transferred to PVDF membrane (BioRad) in transfer buffer (25mM Tris, 190mM glycine and 0.1% SDS) overnight at 4°C at constant voltage (60V). Following the transfer, membranes of SDS-PAGE were washed with 1X TBS with 0.1% Tween-20 (TBST) and blocked with 4% BSA (in TBST) overnight at 4°C. BN-PAGE membranes were first air-dried for 8h at RT and then destained with methanol. After fixation in 5% acetic acid for 30min and extensive washing in TBST, membranes were blocked in 4% BSA (blocking buffer). PVDF membranes were probed with the desired primary (1:1000 dilution) or secondary antibodies (1:10,000 dilution) at 4°C. Non-specific binding was removed by washing the membrane thrice in TBST with 2% BSA for 10 min each. Followed again, by washing with TBST, membranes were developed by Clarity Western ECL substrate (BioRad) and images were acquired by the chemiluminescence gel doc system (Syngene, USA). Native protein standards were cut from the BN-PAGE membranes post-transfer and stained separately with Coomassie G250 to identify the immune-reactive complexes based on the M.W.

### Confocal microscopy and spectral imaging

The spectral images (512×512 pixel resolution, an average of 4 scans) were captured with 4× optical zoom using 60× oil-immersion objective (NA=1.4) under the operation of NIS Elements version 3.0 software. Linear unmixing of the fluorescence spectral images was performed using the PoissonNMF plugin (https://neherlab.org) for the image in Fiji open-source platform (v1.47) (https://imagej.net/Fiji) (41). Although with an option of blind unmixing, the plugin was used to unmix the emission spectra to obtain total donor, acceptor and FRET signals from the reference spectra of the pure fluorophores obtained from the similar instrumental settings. The background was considered as separate spectra and was eliminated by the plugin during unmixing. In our spectral imaging, we eliminated cross-excitation and bleed-through by linear unmixing in order to obtain FRET-based quantities **ξ**^D^ and **ξ**^A^. To obtain **ξ**^D^, unmixed FRET and donor images were added and FRET image was divided by this summed image. The algorithm generated the pixel-wise **ξ**^D^ distribution in images as per the equation:

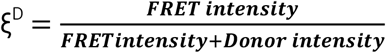

**ξ**^D^ corresponds to FRET efficiency.

Similarly, to obtain the pixel-wise **ξ**^A^ distribution, unmixed FRET images were divided with total acceptor intensity as:

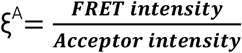

It provides a normalized quantity that represents a fraction of acceptor sensitized by the donor. The implication of this quantity in detecting lower or higher-order p53 homo-oligomeric assemblies is authenticated by CHX-trap based anti-p53 BN-PAGE immune-blotting.

All the pixel-wise calculations were performed through the in-built operations in Fiji. For the random selection of pixels in the **ξ**^D^ and **ξ**^A^ images, add random ROIs (region of interests) plugin in Stowers ImageJ plugins collection (http://research.stowers.org/imagejplugins) was used. A binary mask option in the plugin was used to collect pixel-wise measurements through automatically generated random point ROIs above the threshold and values were used to generate histograms. Friedmann-Diaconis rule was followed while deciding the bins for the histogram of the **ξ**^D^ and **ξ**^A^ distribution. The probability distribution was fitted with one or two- Gaussian functions in spreadsheets (MS-Excel, 2007) using Solver add-in in order to determine the mean values for comparison in different conditions.

In the three-chromophore F-TEPA setup, cross-excitation and bleed-through effects were insignificant under our acquisition settings and therefore required no further corrections.

### Apoptosis analysis

For cell death assay, treated cells were rinsed with PBS and trypsinized. The cells were harvested in the 1mL ice-cold PBS and tapped gently to re-suspend the clumps. Following centrifugation (200× g at 4°C) and aspiration of the supernatant, cells were re-suspended in 1mL ice-cold PBS. The apoptosis was analyzed with the Vybrant Apoptosis Assay Kit as per the manufacturer’s instructions by BD FACS Calibur flow cytometer.

### Plasmids and transfection

ECFP-HIF-1α and EYFP-HIF-1β constructs were kind gifts from J. Fandrey (42). p53-EGFP and p53-DsRed Ex plasmid constructs were generated using the pN1-EGFP and pN1-DsRed Ex vectors (Clontech). All the transfections were performed with Lipofectamine 3000 following the product manual. Briefly, 1.5×10^4^ cells were seeded over the coverslips (gelatine coated) in the wells of 6 well plates 24h prior transfection. A total of 1μg plasmid DNA was used to transfect cells whether single or in co-expression conditions. The complex of the transfection reagent and plasmids were prepared in opti-MEM reduced serum medium (Thermofisher, 31985070). Transfected cells were allowed to express the fluorescent proteins for the 24h before any experiment.

### Immuno-fluorocytostaining

After the treatments, cells were rinsed with PBS and fixed with 4% (v/v) paraformaldehyde at RT for 10 min. The reaction was quenched with 1mM glycine and cells were washed thrice with PBS. The cells were further fixed with chilled 100% methanol for 5min for permeabilization followed by washing with PBS. To remove the background interference due to auto-fluorescence, fixed cells were treated with NaBH_4_ for 20 min at RT (19). Cells were washed with PBS and blocked in 4% BSA (in PBS, 0.1% TX-100) solution overnight at 4°C. anti-p53 DO1(N-ter *specific)*, anti-p53 C-19 (C-ter specific) and A11 primary antibodies (1:1000 dilution in blocking solution, 4h at 22°C) were used to detect their respective epitopes in p53 molecule in vivo in order to generate donor-acceptor FRET pairs by FITC- or TRITC tagged secondary antibodies. After the incubation with primary antibodies, cells were washed with a blocking solution and probed with secondary antibodies (1:5000 dilution in blocking solution) for 2h in dark at RT. From now onwards all the processing was performed under the dark. Non-specific fluorescent secondary antibodies were washed with blocking solution. Depending upon the FRET experiments, the nucleus of the cells was stained with the 170 nM DAPI solution (optimizable) in PBS for 5 min at RT. SlowFade antifade kit was used to mount the coverslips over the glass slides for confocal imaging.

### Luciferase assay

Luciferase assay was performed with Pierce Firefly Luc One-Step Glow Assay Kit as per manufacturer’s instruction. Cells were co-transfected with HRE-Luc and HIF-1α plasmids constructs and effect on the luciferase activity was analyzed by the transfection of p53 siRNA or HIF-1α shRNA in 6 well plates. Cells were harvested for the analysis by the measurement of the normalized luminescence counts due to expressed luciferase. For the normalization, the plasmid construct of β-galactosidase was co-expressed in equal amounts in wells and expression was measured in the presence of the ONPG substrate (Merck, 369-07-3). The reaction was stopped with 1M Tris-Cl (pH 11) and absorbance was detected at 420 nm in each well by visible spectrophotometer (Perkin Elmer).

### FRET on beads assay

HCT116 p53−/− cells co-transfected with ECFP-HIF-1α, EYFP-HIF-1β and p53-DsRed Ex fluorescent constructs in 6 well plates were exposed to hypoxia (1% O_2_) for 48h duration and cells were lysed in native lysis buffer (as described above) except DNase. All the further processing was done in the dark to minimize the bleaching of the fluorophores. The suspension was centrifuged (1000× g, 25°C) for 10 min and supernatant was collected and added with 100 mM KCl, 0.2 mM EDTA, 20% glycerol, 0.25 mg/ml BSA. Biotin-labeled VEGF dsDNA promoter (10nM) (5’CCACAGTGCATACGTGGGCTC3’, 3’GGTGTCACGTATGCACCCGAG5’) (43) was added and incubated overnight at 4°C. The biotin-labeled dsDNA was pulled with 10 μL Pierce Streptavidin Agarose beads (Thermofisher, 20347) at 25°C for 2h. After washing with 100 mM KCl, 0.2 mM EDTA, 20% glycerol, 0.25 mg/ml BSA solution, the beads were mounted over the glass slide (as described for Immunofluorocytostaining of the cells) to perform the acceptor bleaching by the Zeiss LSM 510 Confocal microscope. DsRed Ex was bleached with 100% Transmission of 543 nm laser. ECFP intensity was captured in 475-525 BP filter after excitation with a 457nm laser. For the capture of the EYFP (ex 514 nm) and DsRed Ex (Ex 543nm), BP 530-600 and LP585 filters were used respectively. Similar filter sets were used in three chromophore FRET experiments by sensitized emission methods for ECFP-EYFP-TRITC pair.

### Cell fractionation

Cell fractionation for the analysis of p53 subcellular distribution by SDS-PAGE was performed by the REAP method (44). Briefly, treated cells washed and scrapped in chilled PBS followed by pop-spin for 10 sec in tabletop centrifuge (Eppendorf). After discarding supernatants, cells were resuspended in 900μl chilled 0.1% NP-40 in PBS. After trituration with 1000μl pipette, 300μl suspension was removed as a whole-cell lysate. The remaining suspension was centrifuged for 10 sec at 10,000 rpm in a tabletop centrifuge and 300μl suspension was removed as cytosolic fraction. The remaining supernatant was discarded and nuclear pellet was washed with chilled 0.1% NP40 in PBS. After washing, nuclear pellet was resuspended in 180μl laemmeli buffer which provided nuclear fraction. The nuclear suspension was sonicated followed by boiling in sample loading buffer. Whole-cell, cytoplasmic and nuclear fractions were loaded on gel and SDS-PAGE was performed to analyze p53 subcellular distribution by western blotting.

### Densitometry and processing of western blots

All the densitometry measurements of the HRP luminescence from the immune-blots were performed with Fiji (v1.47) software. The immune-density of the bands was normalized to the density of GAPDH in the same lanes to represent the signal. Normalized signals were used for further calculations. For the proper visualization and representation of immune blot images, lighting adjustments (brightness & contrast or level) were minimally applied over the entire image and performed only for improved blot representation and visualization purposes. All the quantitative measurements were performed with unprocessed 1D BN- or SDS-PAGE immune blot. Due to very low level of endogenous p53T-HIF-1 level captured by 2D BN-PAGE immune blotting, minimal lighting level adjustments were first performed over entire anti-p53 and anti-HIF-1α blot images for proper visualization and representation of dissociated species in the merged image and second, color balance was adjusted over entire merged anti-p53 (magenta) and anti-HIF-1α (cyan) immune blots for the clear visualization of the color changes (purple) upon overlapping.

### Merging immune blots in silico

BN-PAGE anti-p53 DO1 immune blots were stripped with 1M Tris-HCl pH 6.8, 10 ml of 20% SDS and 700 μl β-mercaptoethanol followed by extensive washing in TBST to remove the traces of β-mercaptoethanol. The blots were again blocked with 4% BSA before re-probing with the anti-HIF-1α antibody. A similar stripping procedure and blocking were performed before anti-HIF-1β probing. The HRP luminescence images of both the antibodies were aligned, merged and pseudocolored in Fiji.

### Experiment Materials

Details of materials utilized in the experiments are listed in Table S1.

## Acknowledgments

S Das, HCT116 p53+/+ and HCT116 p53−/− cells; J Fandrey, plasmid constructs of ECFP-HIF-1α and EYFP-HIF-1β; R. Kulshreshtha, HIF-1α shRNA; AIRF, JNU for Confocal imaging; S Bhattarai for p53-DsRed Ex construct; P Malhotra for support in Flow cytometry.

## Author contributions

ST & UP conceived the idea; ST conducted experiments; ST and UP analyzed data and wrote the manuscript.

## FOOTNOTES

Funding was provided by the Indian council of medical research (ICMR), Govt. of India through UPOE grants, as well as Ph.D. Fellowship to S.T. from Council of Scientific and Industrial Research (CSIR), Govt. of India.

The abbreviations used are:

p53: 53kDa tumor protein
BN-PAGE: Blue-native Polyacrylamide gel electrophoresis
D: Dimer
FRET: Förster resonance energy transfer
F-TEPA: FRET-based terminal ends proximity analysis
HIF-1: Hypoxia-inducible factor-1
H.O.: Higher-order
M: Monomer
MDM2: Mouse double minute 2 homolog
ML: mutant-like
O: Octamer
O_2_: Oxygen
R.A.: relative abundance
R.E.: Response element
T: Tetramer
WT: wild type

