## Supplementary Figures and Table for "Oxygen-responsive p53 tetramer-octamer switch controls cell fate"

From the Transcription & Human Biology Laboratory, School of Biotechnology

Authors' addresses

This PDF file includes:

Figs. S1 to S4

Table S1

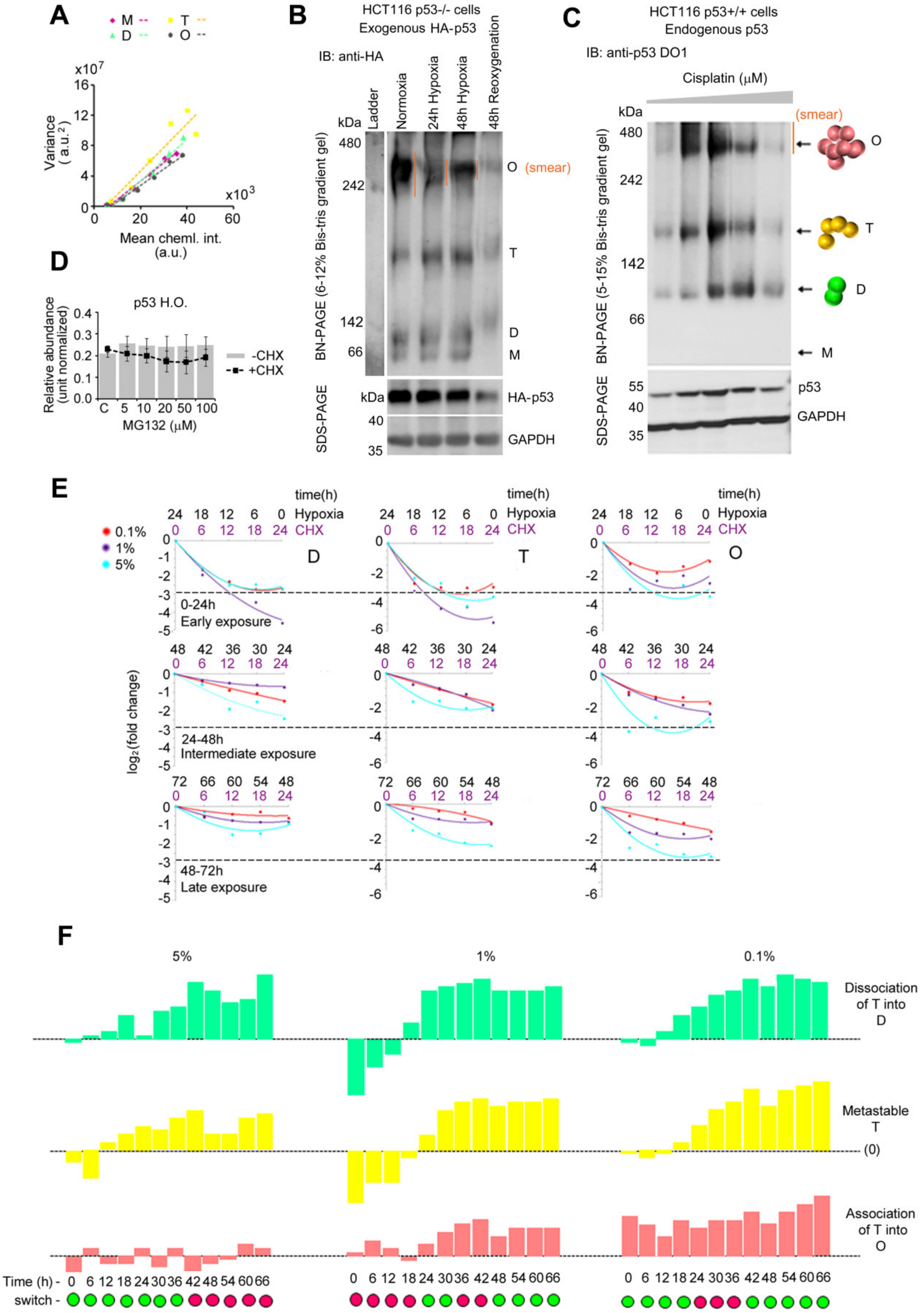

**Figure S1.** (A) Verification of pixel saturation by HRP chemiluminiscence in BN-PAGE immune blots upon extended exposure to camera. Mean v/s variance plot was generated by determining the mean and variance of chemiluminiscence intensity distribution in the pixels of each immune band corresponding to p53 species. Linear regression was performed to check pixel saturation for further densitometry based measurements. (B) An exogenous system consisting of HA-p53 expressed in HCT116 p53-/- cells shows modification of O in the form of smear by anti-HA BN-PAGE immune blot under normoxia (21% $O_2$), hypoxia (1% $O_2$) or re-oxygenation (48h). (C) An endogenous system of HCT116 p53+/+ cell line shows similar O smear by dose-dependent cisplatin treatment (10-50 μM). At higher doses, O smear dissapears with p53 degradation suggesting modification via ubiquitination. (D) R.A. for p53 H.O. determined from -CHX and +CHX variants of trap in the basal state of U2OS cells. (E,F) T-O switching pattern in a gradient of hypoxia.

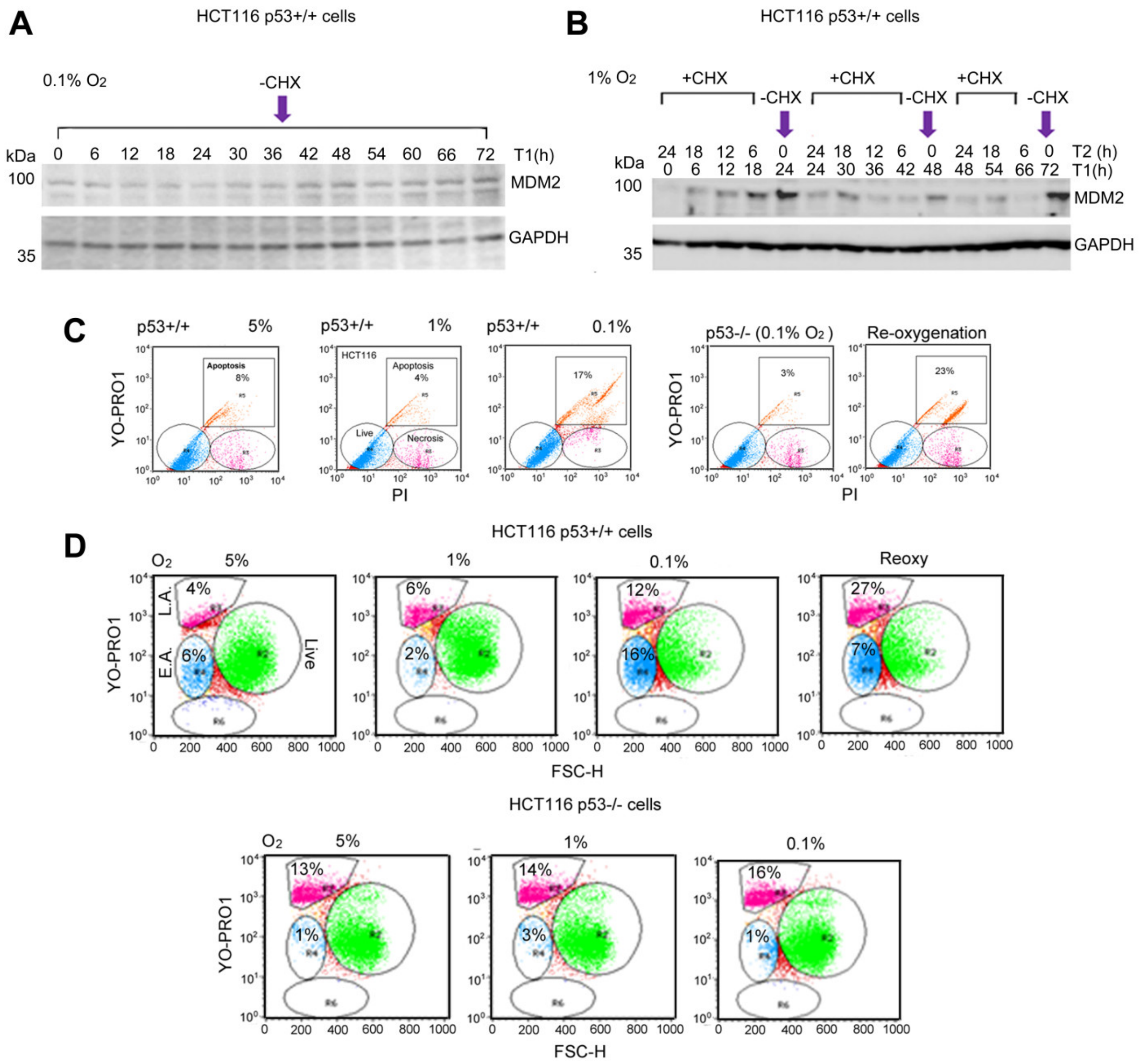

**Figure S2.** SDS-PAGE based MDM2 dynamics without trap at 0.1% $O_2$ (A) and with trap at 1% $O_2$ (B). Immune blots are the best representation of data in Fig. 3B. Apoptosis analysis by YO-PRO1 v/s PI (C) or FSC (forward scatter) v/s YO-PRO1 (D). Analysis in (D) required no compensation during the acquisition of data. 10,000 cells were analyzed and used to generate scattered plot in (C,D) by flow cytometry.

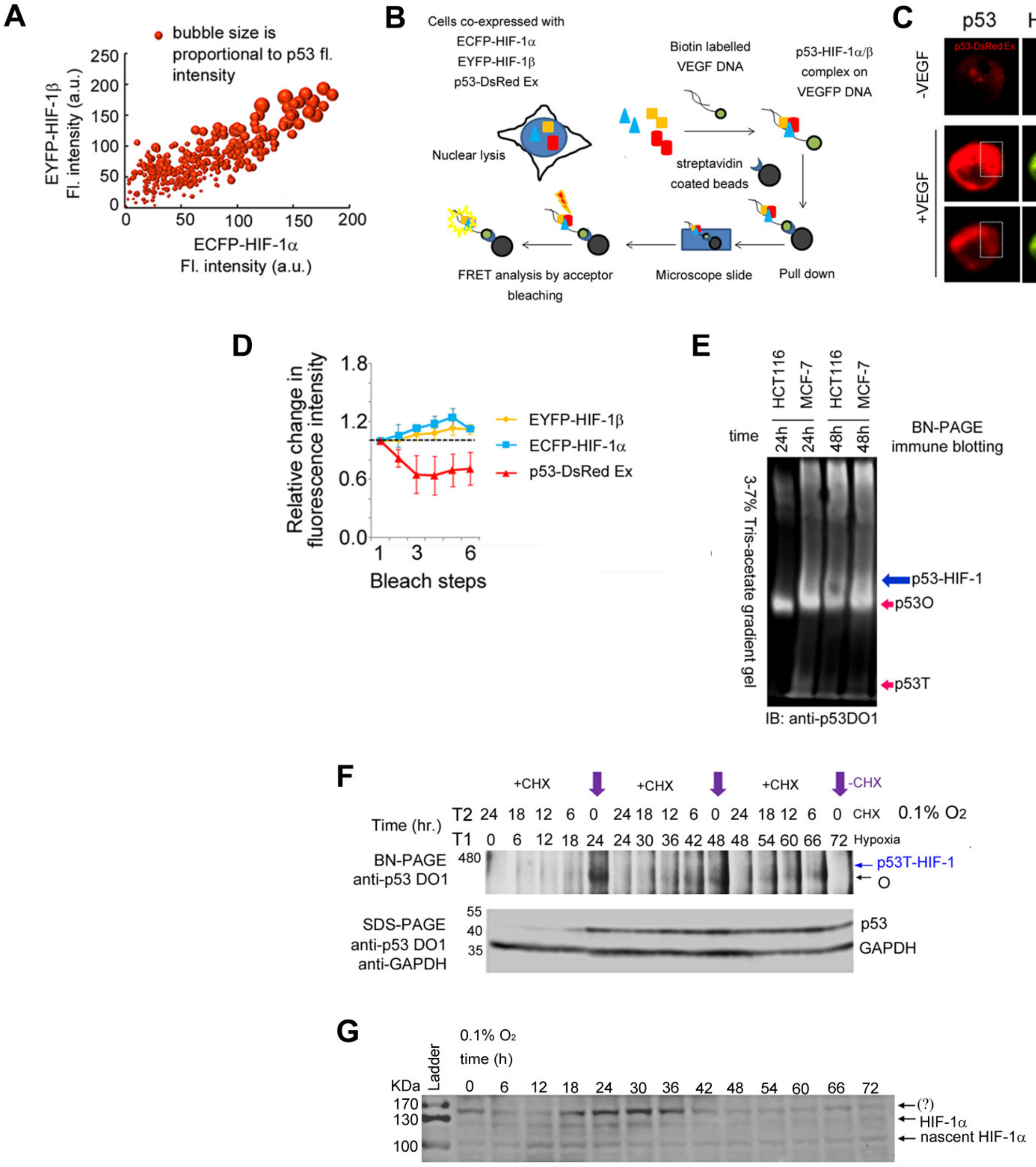

**Figure S3.** (A) Quantification of concentration dependent sequestration of endogenous p53 by exogenous HIF-1 subunits. Bubble size is proportional to the p53 fluorescence intensity. (B,C) FRET on bead assay, a variant of antibody based pull down assays to analyze p53-HIF-1 complex upon VEGF promoter by acceptor bleaching based FRET determination method. (D) p53-HIF-1 complex association by sequential acceptor bleaching inside the cell. (E) segregation of overlapping p53 O and p53-HIF-1 immune bands in Fig 4E by 3-7% Tris-acetate gradient (pH 7.0). (F) p53T-HIF-1 dynamics at 0.1% $O_2$. (G) HIF-1α dynamics at 0.1% $O_2$. The GAPDH loading control for immune blot in (G) is shown in Fig. S2A.

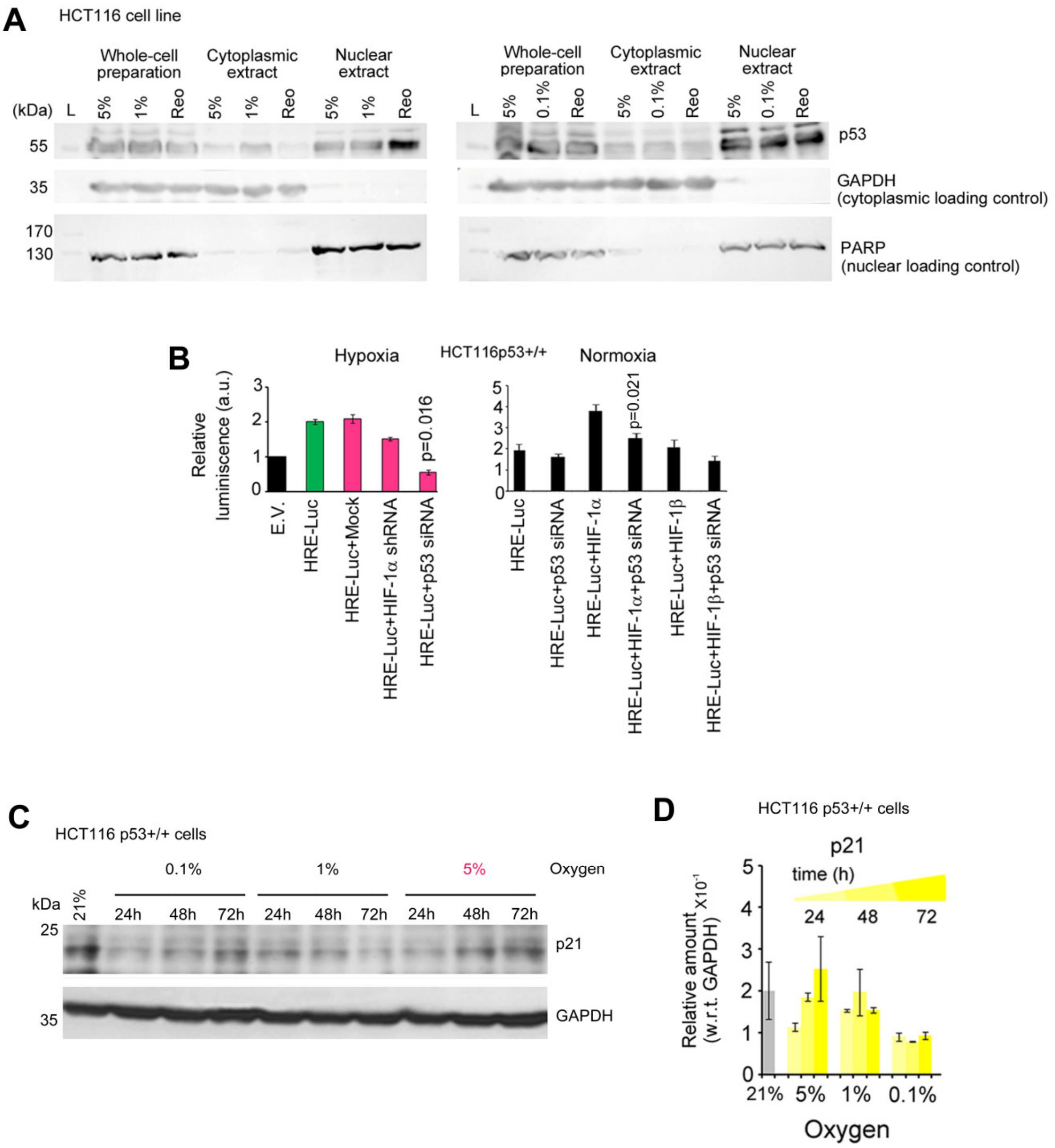

**Figure S4.** (A) SDS-PAGE immune blots shows subcellular localization of p53 in cytoplasmic or nuclear fraction at 1 or 0.1% $O_2$. (B) Luciferase assay shows transcriptional activation of HIF-1 under distal arrangement of its subunits in presence of p53. (C,D) p21 dynamics in hypoxia gradient by SDS-PAGE immune blotting.

| **RESOURCE** | **SUPPLIER** | **IDENTIFICAT** |
|---|---|---|
| **Antibodies** | | |
| Mouse monoclonal p53 (Do1) | Santa Cruz Biotechnology | Cat# Sc-126 |
| Mouse monoclonal HIF-1a (H1alpha67) | Novus Biologicals | Cat# NB100-105 |
| Mouse monoclonal MDM2 (SMP14) | Santa Cruz Biotechnology | Cat# Sc-965 |
| Goat polyclonal p53 (C-19) | Santa Cruz Biotechnology | Cat# Sc-1311 |
| Mouse monoclonal HIF-1beta (H1beta234) | Novus Biologicals | Cat# NB100-124 |
| Rabbit Anti-mouse FITC-tagged secondary | Thermofisher | Cat# 31555 |
| Goat Anti-Mouse TRITC-tagged secondary | Merck | Cat# T7782 |
| Rabbit Anti-Goat TRITC-tagged secondary | Merck | Cat# T7028 |
| Goat Anti-Mouse HRP-tagged secondary | Santa Cruz Biotechnology | Cat# Sc-2005 |
| Rabbit polyclonal anti-oligomer (A11) | Thermofisher | Cat# AHB0052 |
| Mouse monoclonal GAPDH | Santa Cruz Biotechnology | Cat# Sc-47724 |
| Chemicals | | |
| Cisplatin | Merck | Cat# 15663-27-1 |
| Resveratrol | Merck | Cat# 501-36-0 |
| Cycloheximide | Merck | Cat# 66-81-9 |
| Sodium Deoxycholate | Merck | Cat# 302-95-4 |
| CHAPS | Merck | Cat# 331717-45-4 |
| Coomassie G250 | Merck, | Cat# 6104-58-1 |
| Bis-tris | Amresco | Cat# 0715-500G |
| Tricine | Merck | Cat# 5704-04-1 |
| NativeMark protein standards | Invitrogen | Cat# LC0725 |
| MOPS | Merck | Cat# 1132-61-2 |
| Critical Commercial Assays | | |

| | | |
|---|---|---|
| Vybrant Apoptosis assay Kit | Thermofisher | Cat# V13243 |
| SlowFade antifade kit | Thermofisher | Cat# S2828 |
| Pierce Firefly Luc One-Step Glow Assay Kit | Thermofisher | Cat# 16196 |
| Experimental Models: Cell Lines | | |
| HCT116 p53+/+ | (Sen, Satija and Das, 2011; Sen *et al.*, 2013) | NA |
| HCT116 p53-/- | (Sen, Satija and Das, 2011; Sen *et al.*, 2013) | NA |
| U2OS | NCCS, Pune | NA |
| MCF-7 | NCCS, Pune | NA |
| Oligonucleotides | | |
| Biotin-labeled VEGF promoter 5'CCACAGTGCATACGTGGGCTC3', 3'GGTGTCACGTATGCACCCGAG5' | As described by Olenyuk *et al.,* 2004 | NA |
| Recombinant DNA | | |
| ECFP-HIF-1α | Wotzlaw C *et al.*, 2006) | NA |
| EYFP-HIF-1β | Wotzlaw C *et al.*, 2006) | NA |
| HIF-1α shRNA | A gift from Kulshreshtha R. | NA |
| p53 siRNA | Santa Cruz Biotechnology | Cat#sc-29435 |
| Software and Algorithms | | |
| Fiji (image j) version 1.47 | https://imagej.net/Fiji | |
| PoissonNMF | https://neherlab.org | |
| Add random roi plugin | http://research.stowers.org/imagejplugins | |
| Other | | |
| Propidium iodide | Sigma | Cat# P4864 |
| Lipofectamine 3000 | Invitrogen | Cat# L3000015 |

| | | |
|---|---|---|
| DAPI | Thermofisher | Cat# D1306 |
| PVDF membrane | BioRad | Cat# 1620177 |
| Clarity western ECL substrate | BioRad | Cat# 1705060 |
| Hypoxia Chamber | Plas Labs inc. | Model# 856-HYPO |

**Table S1.** Details of materials used in experiments.
